# NLRP3 inflammasome signaling orchestrates the cellular and spatial architecture of hepatic granuloma and protective immunity

**DOI:** 10.64898/2026.09.18.752760

**Authors:** Isadora M. de Oliveira, Mariana M. Chaves, Juliana Costa-Madeira, Ana Luisa Barbosa, Pedro S. Corradi, Bianca de Oliveira, Amanda M. Becerra, Pedro Henrique Marques, Tamara S. Rodrigues, Adriana Simizo, Lucas Lorenzon, Sabrina S. Batah, Andrea J.R. Herrera, Helder Nakaya, Amanda M. da Silva, Ricardo T. Gazzinelli, Angela K. Cruz, Cristina M. Takiya, Alexandre T. Fabro, Gustavo B. Menezes, Carlos H.N. Costa, Dario S. Zamboni

## Abstract

Granulomas are organized immune structures that contribute to host defense against persistent pathogens, yet the mechanisms that coordinate their assembly and protective function remain incompletely understood. Here, using experimental visceral leishmaniasis caused by *Leishmania infantum* and samples from patients with active disease, we identify the NLRP3 inflammasome as a regulator of protective hepatic granulomatous immunity. Inflammasome-associated mediators were elevated in patients and correlated with systemic inflammation. In mice, *L. infantum* induced NLRP3 inflammasome activation within hepatic granulomas, while single-cell and spatial transcriptomic analyses revealed enrichment of inflammasome-associated transcription in hepatic macrophages and granuloma-associated regions. NLRP3 deficiency did not prevent granuloma initiation but impaired granuloma expansion, cellular organization, and leukocyte accumulation. This response required Caspase-1/11 and IL-18, but was independent of IL-1β. Loss of NLRP3 also impaired parasite control despite reducing hepatic inflammation and histopathological alterations. Together, our findings identify NLRP3-Caspase-1/11-IL-18 signaling as a mechanism that coordinates the cellular and spatial organization of protective hepatic granulomas, revealing granuloma architecture as a previously unrecognized function of inflammasome-mediated immunity and providing mechanistic insight into host resistance in visceral leishmaniasis, a potentially fatal neglected tropical disease.

## Introduction

Granulomas are organized multicellular inflammatory structures that arise in response to persistent infectious or non-infectious stimuli. Rather than representing simple cellular aggregates, granulomas are composed of spatially arranged immune cell populations, including macrophages and lymphocytes, whose organization can influence pathogen containment, tissue inflammation, and disease outcome (Pagan and Ramakrishnan, 2018). In infectious diseases, granuloma formation has been classically associated with host defense by restricting pathogen dissemination, although these structures may also provide niches for pathogen persistence and contribute to tissue pathology (Cohen et al., 2022; Davis and Ramakrishnan, 2009). Thus, understanding how granulomas are assembled and organized is essential to define how protective granulomatous immunity is established.

Granuloma formation is a highly coordinated process that depends on the recruitment, retention, and spatial organization of multiple immune cell populations. Studies in tuberculosis have shown that the composition and architecture of granulomas evolve over time and are closely associated with their ability to contain pathogens (Davis and Ramakrishnan, 2009; Guirado et al., 2013; Marakalala et al., 2016). Chemokines, cytokines, and cellular interactions contribute to the assembly of these structures by regulating leukocyte trafficking and positioning within affected tissues (Cronan et al., 2016; Marakalala et al., 2016; Pagan and Ramakrishnan, 2018). However, despite substantial advances in defining the cellular composition of granulomas, the mechanisms that coordinate their structural organization and functional maturation remain incompletely understood.

Innate immune sensing pathways are among the earliest regulators of tissue inflammation and therefore represent attractive candidates for controlling granuloma assembly. Among these, inflammasomes are multiprotein complexes that detect microbial products and cellular stress signals, leading to activation of inflammatory caspases and the maturation of IL-1 family cytokines (Broz and Dixit, 2016; Schroder and Tschopp, 2010). In addition to their established roles in antimicrobial defense, inflammasomes have emerged as important regulators of leukocyte recruitment, cellular activation, and inflammatory tissue responses in a variety of settings (Swanson et al., 2019).

These observations raise the possibility that inflammasome signaling may contribute not only to the intensity of inflammatory responses but also to the organization of immune cells within inflamed tissues. Consistent with this idea, inflammasome-associated pathways have been implicated in several granulomatous disorders, including sarcoidosis and granulomatous alveolitis (Denis, 1994; Huppertz et al., 2020). Nevertheless, it remains unclear whether inflammasomes simply amplify inflammatory responses within granulomas or actively regulate the assembly, structural organization, and function of these structures as protective immune niches.

Visceral leishmaniasis provides a valuable model to investigate the relationship between inflammation, granuloma organization, and host protection. In both experimental and human disease, visceralizing *Leishmania* species induce the formation of hepatic granulomas that are critical for parasite control (Engwerda et al., 1998; Kaye and Beattie, 2016; Lodi et al., 2024; Murray, 2001; Murray et al., 1992; Stanley and Engwerda, 2007). Studies in experimental visceral leishmaniasis have shown that granulomas undergo a progressive maturation process characterized by the recruitment of distinct immune cell populations, and that the acquisition of an organized granulomatous architecture is closely associated with the development of resistance (Engwerda et al., 2004; Murray, 2001). Consequently, this model offers a unique opportunity to investigate how innate inflammatory pathways influence the assembly and function of protective granulomatous responses.

Based on the central role of hepatic granulomas in resistance to visceral leishmaniasis and the established functions of inflammasome signaling in inflammatory tissue responses, we hypothesized that inflammasomes contribute to host protection not only by promoting inflammation but also by regulating the organization of protective granulomatous immunity. To address this question, we first evaluated evidence of inflammasome activation in patients with active visceral leishmaniasis and subsequently established a murine model of *Leishmania infantum* infection to investigate the contribution of NLRP3 inflammasome signaling to granuloma formation, cellular organization, and host resistance. Our findings provide evidence of systemic inflammasome activation in patients with visceral leishmaniasis, establish a protective role for NLRP3 inflammasome signaling in experimental disease, and uncover a previously unrecognized function for inflammasomes in coordinating the organization of protective granulomatous immunity.

## Results

### Inflammasome-associated mediators are increased and correlate with systemic inflammatory responses in patients with visceral leishmaniasis

Although genetic studies and experimental models have implicated IL-18 and inflammasome-associated pathways in host responses to *Leishmania* infection (Ahmadpour et al., 2016; Bhattacharya et al., 2024; Haeberlein et al., 2010; Kumar et al., 2014; Lima-Junior et al., 2013; Moravej et al., 2013; Mullen et al., 2006; Murray et al., 2006; Vieira et al., 2024), whether these pathways are activated during human visceral leishmaniasis remains unclear. To address this question, we analyzed serum samples from patients with active visceral leishmaniasis. Serum samples from 100 visceral leishmaniasis patients were obtained from the Natan Portela Institute of Tropical Diseases (IDTNP), located in Teresina, Brazil, an endemic area for *L. infantum*. Cytokine concentrations were quantified using a CBA assay. Compared with healthy controls, patients with active visceral leishmaniasis displayed significantly increased serum levels of IL-1β (**Figure 1A**), IL-18 (**Figure 1B**), and caspase-1 p20 (**Figure 1C**). These findings indicate increased systemic levels of inflammasome-associated mediators during active disease. To investigate whether these mediators were associated with the inflammatory profile observed in visceral leishmaniasis, we performed Pearson correlation analyses using IL-1β, IL-18, and caspase-1 p20 levels. We observed strong positive correlations between IL-1β and the inflammatory cytokines IL-6, IL-8, IL-12, TNF-α, IL-10, and IFN-γ (**Figure 1D-I**). Similarly, IL-18 levels positively correlated with IL-6, IL-8, and IL-10 (**Figure 1J-O**). In contrast, no significant correlations were observed between caspase-1 p20 levels and any of the cytokines analyzed (**Figure 1P-U**). Collectively, these findings demonstrate that inflammasome-associated mediators are elevated during active visceral leishmaniasis and are closely linked to systemic inflammatory responses, prompting us to establish an experimental model of *L. infantum* infection to mechanistically investigate the role of inflammasome signaling and its associated cytokines during disease development.

**Figure 1.**
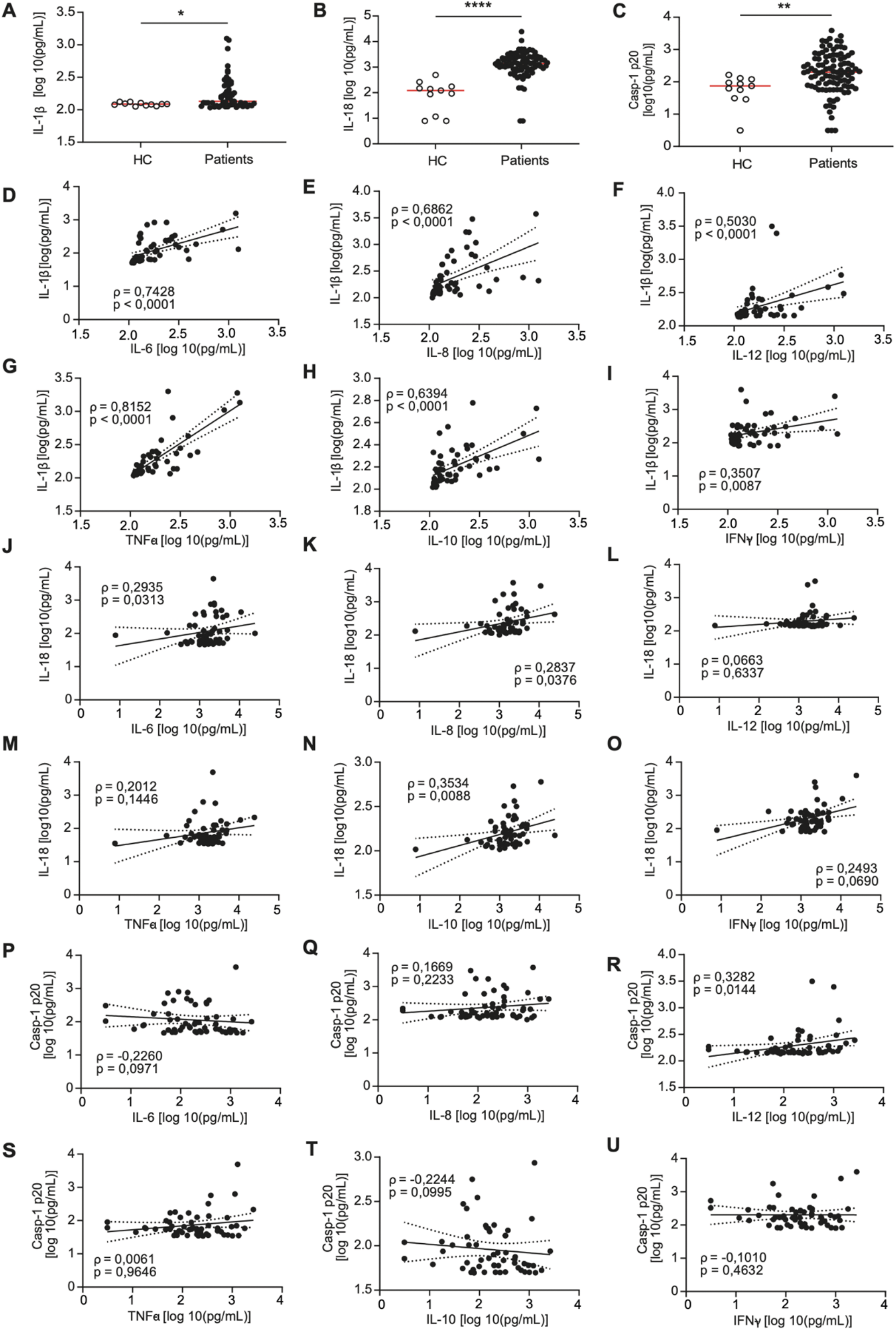
Patients with visceral leishmaniasis exhibit increased circulating IL-1β, IL-18, and caspase-1 p20 associated with systemic inflammatory responses. Serum of patients with VL were collected and the levels of cytokines were analyses by Cytometric Bead Array (CBA) assay. Levels of IL-1β (A), IL-18 (B) and Caspase-1 p-20 (C) were accessed. Pearson’s correlations between IL-1β and IL-6 (D), IL-8 (E), IL-12 (F), TNF-α (G), IL-10 (H) and INF-γ (I). Pearson’s correlations between IL-18 and IL-6 (J), IL-8 (K), IL-12 (L), TNF-α (M), IL-10 (N) and INF-γ (O). Pearson’s correlations between Caspase-1 p20 and IL-6 (P), IL-8 (Q), IL-12 (R), TNF-α (S), IL-10 (T) and INF-γ (U). *, *P* <0.05; **, *P* <0.005; ***, *P* <0.0005; ****, *P* <0.00005.

### Inflammasome-associated transcription is enriched in hepatic macrophages and spatially associated with granulomas

Single-cell and spatial transcriptomic datasets generated from the livers of naïve and *L. donovani*-infected mice (Dey et al., 2026), were reanalyzed to define the cellular and spatial distribution of inflammasome-associated transcripts (**Figure 2A**). The scRNA-seq dataset identified two major hepatic macrophage populations: Lyz2hi_MoMac cells, corresponding to monocyte-derived macrophages, and ApoeHi_Kupffer cells, corresponding to resident Kupffer cells. A gene signature comprising *Nlrp3*, *Pycard*, *Casp1*, *Il1b*, *Il18*, and *Gsdmd* was increased in cells from infected livers and was most prominent within myeloid populations (**Figure 2B**). Pseudobulk analysis comparing hepatic macrophages with all remaining liver cell populations further showed enrichment of all six inflammasome-associated transcripts in the macrophage compartment. The largest differences were observed for *Nlrp3*, *Il1b*, and *Il18*, which exhibited 15.8-, 4.9-, and 22.3-fold higher expression, respectively (**Figure 2C** and **Figure S2A**). Together, these analyses identify hepatic macrophages as a major cellular compartment enriched for inflammasome-associated transcripts in the infected liver.

**Figure 2.**
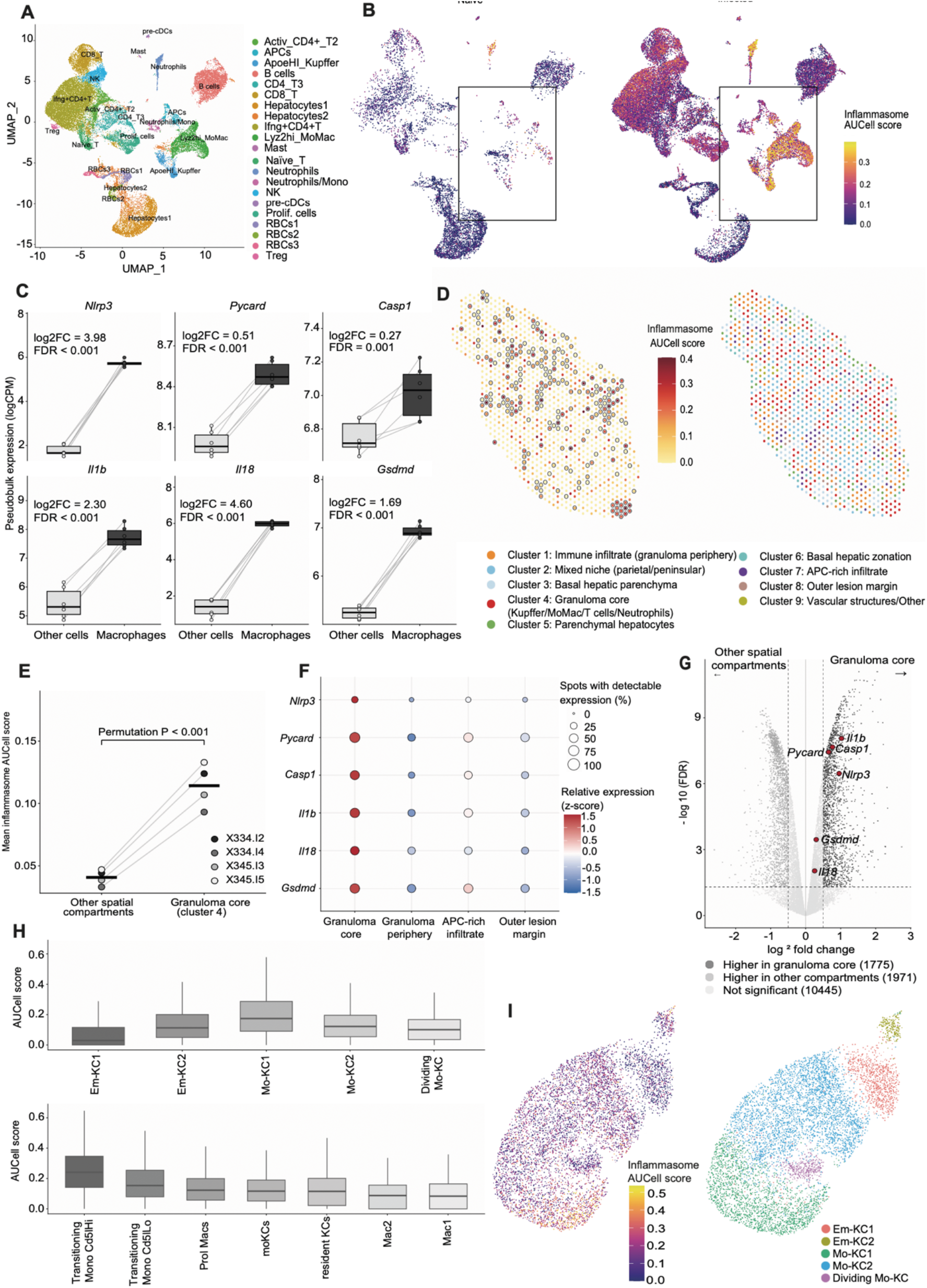
Inflammasome-associated transcription is enriched in hepatic macrophages and spatially associated with granulomas during visceral leishmaniasis. UMAP representation of the single-cell RNA-sequencing dataset from the livers of naïve or infected mice, colored according to the annotated cell populations (A). Distribution of the AUCell score for an inflammasome signature. The boxed regions correspond to the hepatic macrophage compartments (B). Pseudobulk expression, shown as logCPM, of the signature components in macrophages and all other cell populations. Each point represents one biological sample (C). Spatial distribution of the inflammasome AUCell score, left, and annotated transcriptional regions, right, in sample X334.I2. Spots outlined in the AUCell map belong to cluster 4, corresponding to granuloma-associated RNA_4 region (D). Paired comparison of the mean AUCell score between the granuloma-associated RNA_4 region, and the remaining spatial compartments across the four samples analyzed. The *P* value was determined using a permutation test (E). Dot plot showing the expression of the signature components in the granuloma core, granuloma periphery, antigen-presenting-cell-enriched infiltrate and outer lesion margin (F). Volcano plot of differential expression between the granuloma-associated RNA_4 region and the remaining spatial compartments. Components of the inflammasome signature are highlighted in red (G). Distribution of the inflammasome AUCell score across hepatic macrophage states identified in two independent single-cell RNA-sequencing datasets. The upper panel shows embryonically derived Kupffer cells, Em-KC1 and Em-KC2; monocyte-derived Kupffer cells, Mo-KC1 and Mo-KC2; and dividing Mo-KC cells. The lower panel shows transitional Cd5l^Hi^ and Cd5l^Lo^ monocytes, proliferating macrophages, monocyte-derived Kupffer cells, resident Kupffer cells, and the Mac1 and Mac2 populations (H). UMAP representation of the hepatic macrophage dataset, colored by the inflammasome AUCell score, left, and by the annotated cell populations, right. In C and G, FDR values correspond to *P* values adjusted using the Benjamini–Hochberg method.

The spatial distribution of the inflammasome-associated gene set was subsequently examined in liver sections from *L. donovani*-infected mice. The highest gene enrichment score were observed in cluster 4, the transcriptomic cluster that showed the strongest spatial overlap with histologically defined granulomas and was predicted to be enriched in macrophages, CD4^+^ and CD8^+^ T cells, NK cells, and neutrophils (**Figure 2D** and **Figure S1**). Elevated scores within cluster 4 were consistently observed across all four infected spatial samples and were supported by within-sample permutation analyses comparing cluster 4 with the remaining hepatic regions (**Figure 2E**). Gene-level analysis indicated that this enrichment reflected increased expression of individual components of the six-gene signature, although the magnitude of enrichment varied among transcripts. Other immune-enriched spatial regions displayed lower or intermediate expression relative to cluster 4 (**Figure 2F** and **Figure S2B**). Differential expression analysis comparing cluster 4 with the remaining spatial transcriptomic clusters identified 1,757 enriched transcripts, including *Nlrp3*, *Pycard*, *Il1b*, *Il18*, and *Gsdmd* (**Figure 2G**). Thus, granuloma-enriched regions are spatially associated with increased expression of inflammasome-related transcripts. Because individual Visium spots contain multiple cell populations, these analyses establish the spatial association of the signature with granulomatous regions but do not assign its expression to a specific cell type.

The distribution of the same six-gene inflammasome-associated signature was also assessed in an independent scRNA-seq dataset of hepatic macrophages isolated from mice infected with *L. infantum* (Pessenda et al., 2025). Gene enrichment scores varied across macrophage states, with the highest values observed in Mo-KC1 cells, the lowest in Em-KC1 cells, and intermediate values in Em-KC2, Mo-KC2, and proliferating monocyte-derived macrophages (**Figure 2H, I** and **Figure S2C**). At higher annotation resolution, the strongest scores were observed in transitioning monocyte populations, particularly Transitioning Mono Cd5lHi cells, followed by Transitioning Mono Cd5lLo cells. Resident Kupffer cells, more differentiated monocyte-derived Kupffer cell states, and the remaining macrophage populations displayed comparatively lower scores. Thus, this independent dataset reveals substantial heterogeneity in inflammasome-associated transcript abundance within the hepatic macrophage compartment, with the highest transcriptional signature observed in monocyte populations annotated as transitional states. Together, analyses of the *L. donovani* and *L. infantum* datasets identify hepatic macrophages as an important cellular compartment enriched for inflammasome-associated transcripts and further indicate that this signature varies markedly across macrophage states, while spatial transcriptomic analysis associates increased expression of these genes with granuloma-enriched regions.

### Intraperitoneal infection with *L. infantum* establishes a self-limited visceral infection characterized by hepatic granuloma formation

To establish an experimental model suitable for investigating the role of the NLRP3 inflammasome during visceral leishmaniasis, we first compared different visceralizing *Leishmania* strains, including NLC and PP75 of *L. infantum* and LV9 of *L. donovani*, a well-established strain that has been extensively used in murine models of visceral leishmaniasis due to its ability to establish consistent visceral infection. Mice were infected via the intraperitoneal route, a technically simple and reproducible approach that allows rapid and consistent parasite delivery. Among the strains tested, the NLC strain of *L. infantum* most efficiently established visceral infection (**Figure S3A**). Next, we compared infections using stationary-phase promastigotes (SPh) or metacyclic-enriched cultures (MEC), which contain higher proportions of infective metacyclic forms than conventional stationary-phase cultures (Melo et al., 2017). Both parasite preparations established visceral infection when 10^7^ parasites were inoculated intraperitoneally, although MEC parasites induced higher parasite burdens (**Figure S3B**). Furthermore, infection with 10^6^ MEC parasites resulted in detectable parasite loads in both liver and spleen, whereas the same inoculum of SPh parasites failed to establish infection (**Figure S3B**). Together, these results demonstrate that the NLC strain of *L. infantum* efficiently induces visceral infection in mice and that MEC parasites are more effective than SPh parasites at establishing infection, consistent with previous reports highlighting the enhanced infectivity of metacyclic promastigotes (Bates, 2018).

To characterize the kinetics of *L. infantum* infection in this model, C57BL/6 mice were infected intraperitoneally with metacyclic-enriched cultures (MEC) of the NLC strain and monitored over time. Parasites were readily detected in the liver as early as 2 weeks post-infection, reaching peak levels between weeks 3 and 4. Thereafter, parasite burdens progressively declined and became undetectable by week 8, indicating effective control of hepatic infection (**Figure 3A-B**). To more precisely define the peak of infection, parasite loads were quantified weekly following infection. Parasite numbers increased during the first weeks of infection, peaked at weeks 3–4, and subsequently declined (**Figure 3B**). A similar kinetic profile was observed in the spleen, where parasite burdens also reached maximal levels around week 3 and progressively decreased thereafter, with only low numbers of parasites remaining detectable at week 8 (**Figure S4A-B**).

**Figure 3.**
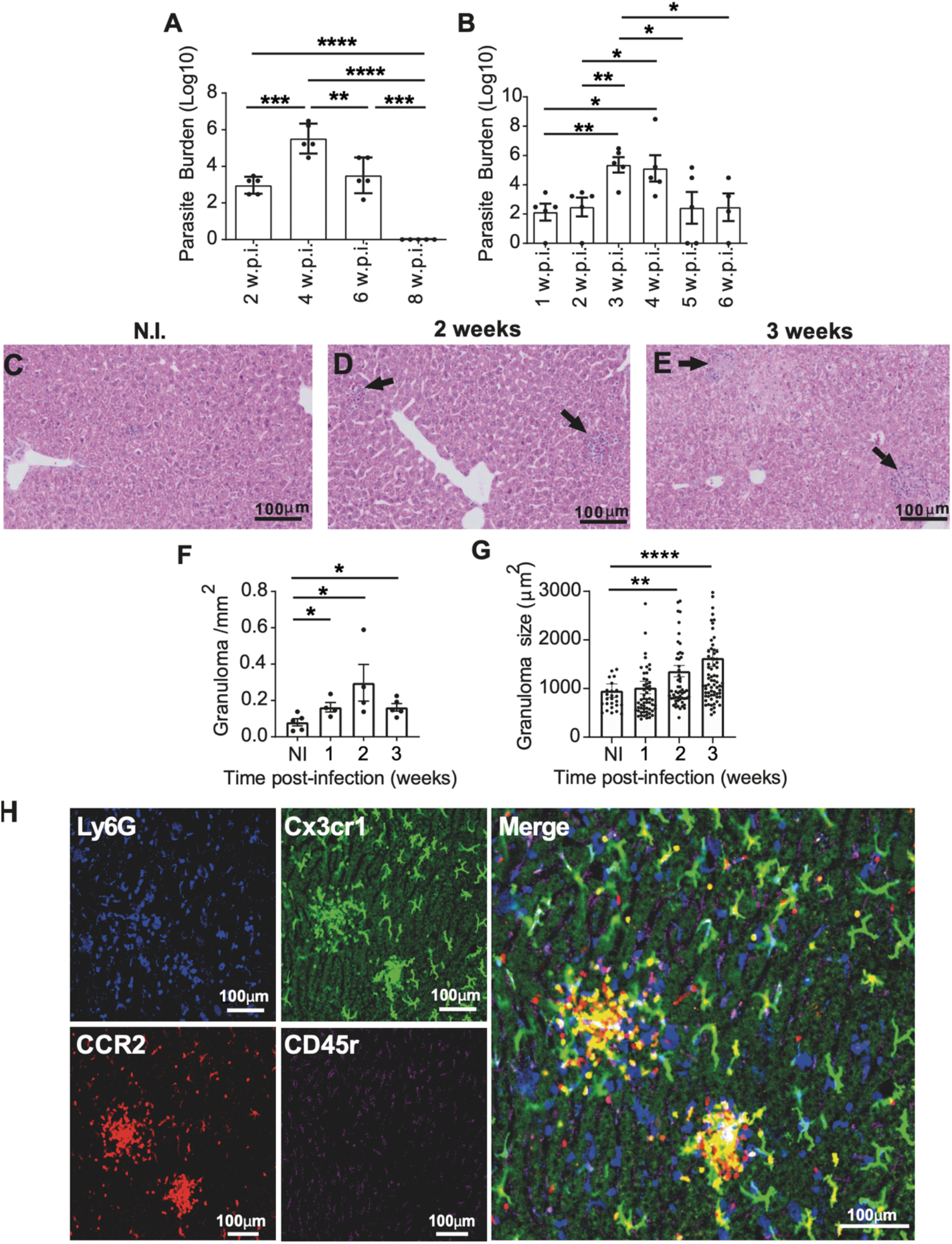
The NLC strain of *Leishmania infantum* establishes a self-limited hepatic infection associated with granuloma development. 10^7^ NLC *L. infantum* were purified from MEC and infected into C57BL/6 mice by IP route and followed for several weeks. After 2, 4, 6 and 8 weeks after the infection, a group of animals (n = 5) were sacrificed and the livers (A) were removed to quantify parasite burden. Also, after 1, 2, 3, 4, 5 and 6 weeks after the infection, a group of animals (n = 5) were sacrificed and the livers (B) were removed to quantify parasite burden. After 2 and 3 weeks of infection, a group of animals (n = 5) were sacrificed and the livers were removed to H&E coloration. Liver from C57BL/6 non-infected (C) and NLC *L. infantum*-infected mice in 2 w.p.i. (D) and 3 w.p.i. (E). The number of granulomas were quantified using Image J (F). Granuloma size were quantified using Image J. Each dot represents a granuloma from a total of 4 or 5 animals (G). One of three independent experiments is shown. Representative image of C57BL/6 Ccr2RFP Cx3cr1GFP animal after three weeks of infection with *L. infantum* NLC (H). Data represented as mean ± DPM; *, *P* <0.05; **, *P* <0.005; ***, *P* <0.0005.

It has been well established that the formation of structurally organized hepatic granulomas is a critical component of acquired resistance to visceral leishmaniasis caused by *L. donovani* (Murray, 2001). Given that parasite burdens peaked around week 3 in our model, we next investigated whether *L. infantum* infection induced granuloma formation during this period. Histopathological analyses revealed a marked recruitment and organization of inflammatory cells within the liver, resulting in the formation of well-defined granulomatous structures consistent with granuloma. Importantly, infected animals displayed significant increases in both the number and size of hepatic granulomas at weeks 2 and 3 post-infection (**Figure 3C-G**).

To further characterize the cellular composition of these structures, we performed intravital microscopy using C57BL/6 Ccr2RFP Cx3cr1GFP reporter mice combined with anti-Ly6G and anti-CD45r staining. Granulomas were composed of CCR2+ cells, including predominantly monocytes and other inflammatory leukocytes, CX3CR1+ cells, Ly6G+ cells, and CD45r+ cells (**Figure 3H**), demonstrating that these structures contain a complex network of myeloid and lymphoid populations. Together, these findings show that intraperitoneal infection of C57BL/6 mice with the NLC strain of *L. infantum* establishes a self-limited visceral infection characterized by peak parasite burdens at weeks 2-3 and the concomitant formation of organized hepatic granulomas.

### NLRP3 inflammasome promotes hepatic granuloma organization through a Caspase-1/11 - IL-18 pathway

Previous studies have demonstrated that IL-1 receptor signaling and NLRP3 inflammasome activation contribute to granuloma formation in inflammatory lung diseases, including alveolitis and sarcoidosis (Denis, 1994; Huppertz et al., 2020). Given the prominent granulomatous response observed during *L. infantum* infection, we next investigated whether NLRP3 contributes to hepatic granuloma formation in this model. Histopathological analyses revealed that *Nlrp3^—/—^*mice infected intraperitoneally with *L. infantum* developed significantly smaller granulomas than infected C57BL/6 mice at 2 weeks post-infection (**Figure 4A-E**). In contrast, although infection increased the total number of granulomas per liver area, no differences were observed between wild-type and *Nlrp3^—/—^* animals (**Figure 4E**). Additional analyses performed at later time points confirmed that NLRP3 deficiency consistently impaired granuloma development and organization at weeks 2, 3, and 4 post-infection (**Figure S5**). These findings indicate that NLRP3 is dispensable for granuloma initiation but contributes to granuloma expansion and structural organization.

**Figure 4.**
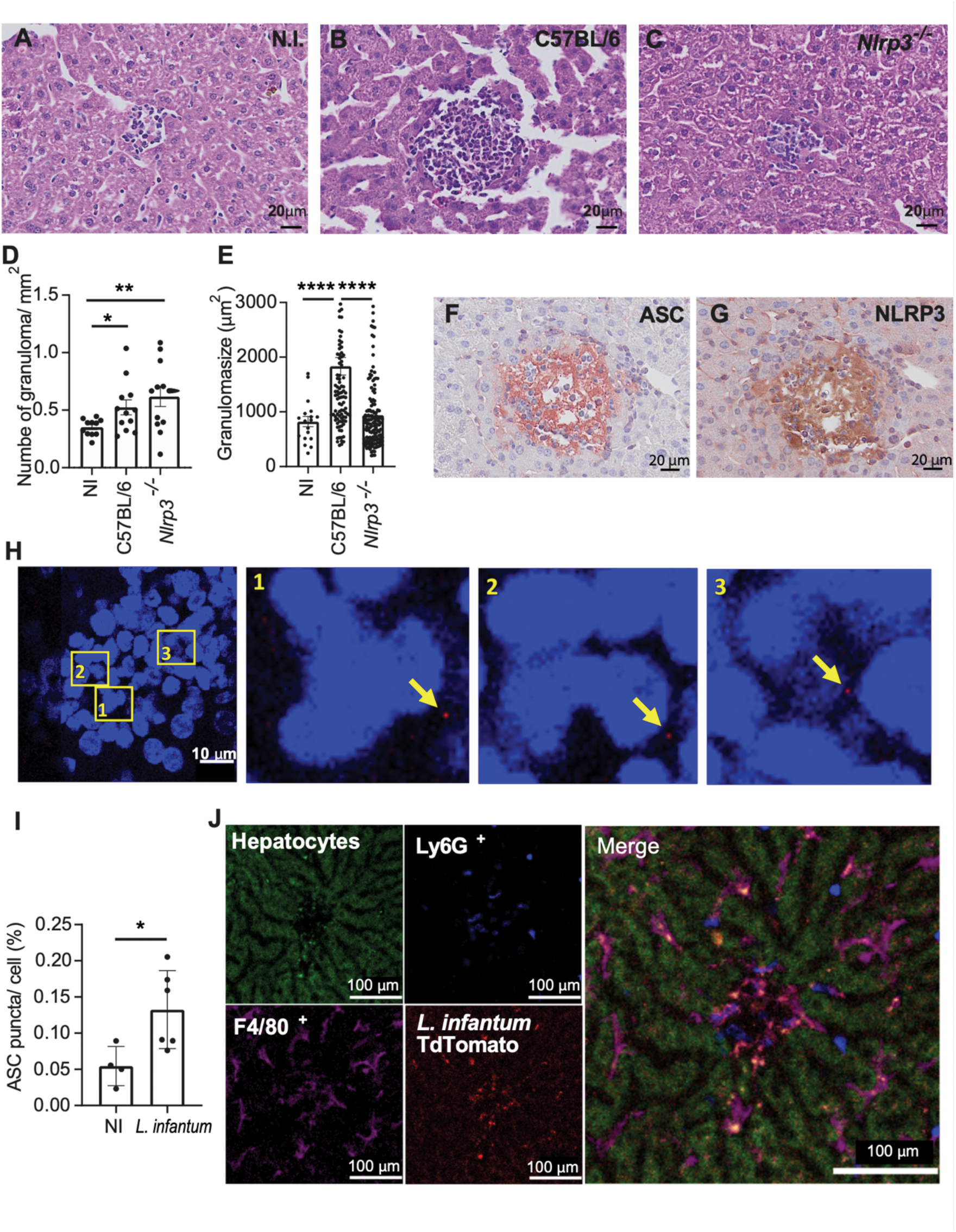
The NLRP3 inflammasome is important for liver granuloma organization in *L. infantum*-infected mice. C57BL/6 and *Nlrp3^—/—^* mice were infected with 10^7^ MEC of *L. infantum* NLC strain via IP, the animals were euthanized and the livers were used for histological analysis. Livers from uninfected (A) and infected C57BL/6 (B) and *Nlrp3^—/—^* mice (C) were obtained after 2 weeks of infection. The number of granulomas were quantified using Image J (D). Granuloma size were quantified using Image J. Each dot represents a granuloma from a total of 11(NI) or 12 (C57BL/6 and *Nlrp3^—/—^*) animals (E). Set of 2 independent experiments were plotted together. C57BL/6 mice were infected with 10^7^ MEC of *L. infantum* NLC strain. Livers from infected mice were collected after 2 weeks of infection and stained for ASC (F) and NLRP3 (G). Representative immunohistochemical images of granulomas in liver tissues. Representative immunofluorescence images of ASC puncta in liver tissues of C57BL/6 mice infected after 2 weeks of infection with 10^7^ *L. infantum* (H). The number of puncta in granulomas was quantified (I). Representative images of granuloma from C57BL/6 animals infected with *L. infantum* NLC TdTomato (J) *, *P* <0.05; **, *P* <0.005; ***, *P* <0.0005; ****, *P* <0.00005.

To determine whether inflammasome components were expressed within these structures, we performed immunohistochemical analyses of infected livers. Both NLRP3 and ASC were readily detected within hepatic granulomas (**Figure 4F-G**), and quantitative analyses demonstrated increased hepatic expression of ASC (**Figure S6A-D**) and NLRP3 (**Figure S6E-H**) following infection. To further investigate inflammasome activation, we assessed ASC speck formation by immunofluorescence. We detected abundant ASC puncta within hepatic granulomas of infected animals (**Figure 4H**), and quantitative analyses confirmed a significant increase in ASC speck formation following infection (**Figure 4I**). Similarly, the total number of NLRP3 puncta in the liver was increased in infected mice (**Figure S6I-L**). Together, these results demonstrate that *L. infantum* infection induces local expression and activation of the NLRP3 inflammasome within hepatic granulomas and that NLRP3 signaling is required for the proper development and organization of these structures.

Having established that NLRP3 contributes to hepatic granuloma development during *L. infantum* infection, we next sought to identify the downstream inflammasome-associated pathways responsible for this phenotype. Since NLRP3 activation promotes caspase-1 activation and the maturation of IL-1β and IL-18, we evaluated granuloma formation in *Casp1/11^—/—^*, *Il1b^—/—^* and *Il18^—/—^* mice. Similar to *Nlrp3^—/—^*animals, infected *Casp1/11^—/—^* and *Il18^—/—^* mice developed significantly smaller granulomas than infected C57BL/6 mice (**Figure 5A-G**). In contrast, granuloma size was not altered in *Il1b^—/—^* mice (**Figure 5A-H**). These findings indicate that the effect of NLRP3 on granuloma development is dependent on the inflammasome pathway and is primarily associated with IL-18 rather than IL-1β. Collectively, these results further support a critical role for the NLRP3 inflammasome in the formation and maturation of hepatic granulomas during *L. infantum* infection.

**Figure 5.**
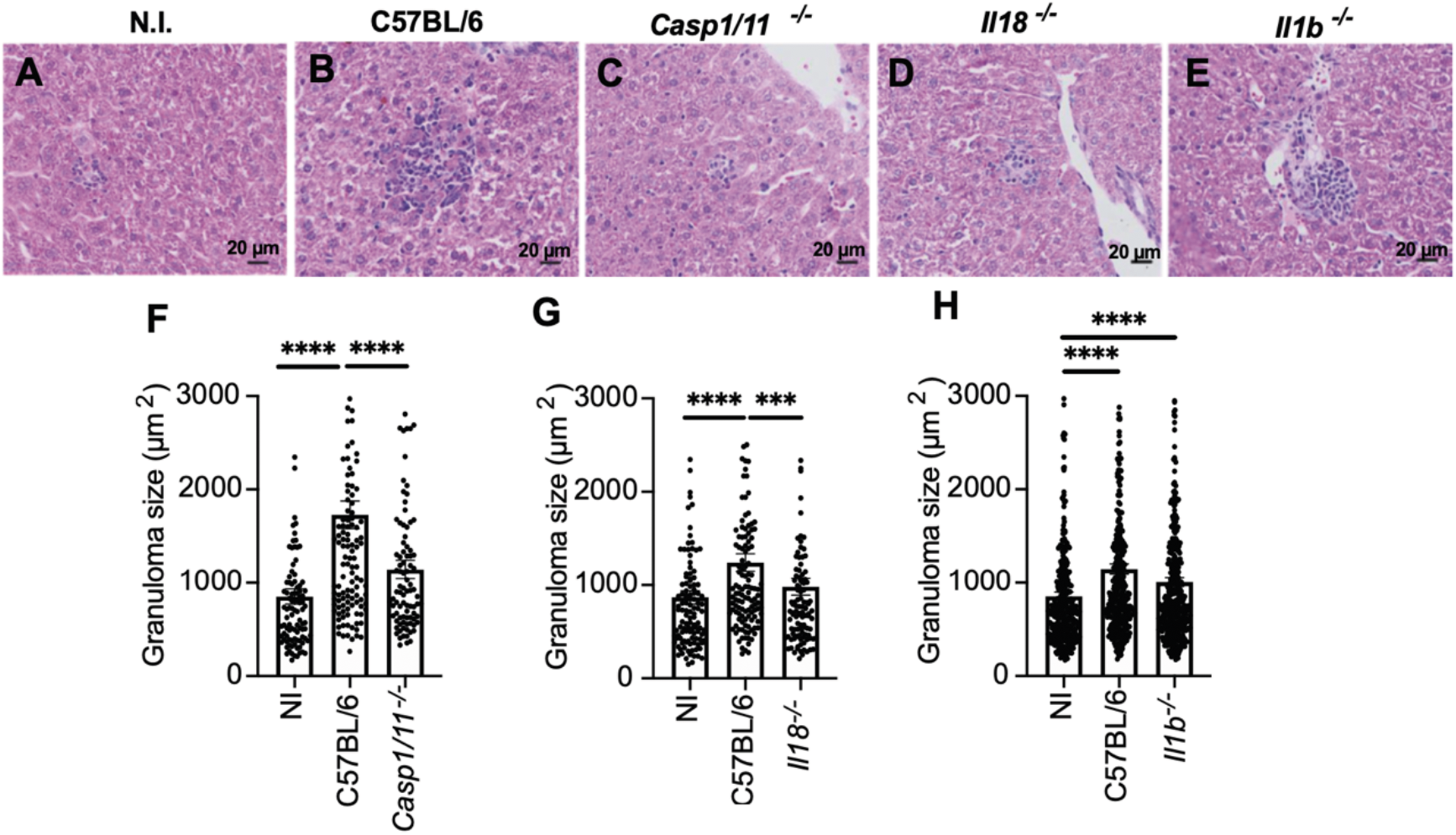
IL-18 mediates NLRP3 inflammasome-dependent hepatic granuloma organization. C57BL/6, *Casp1/11^—/—^*, *Il18^—/—^* and *Il1b^—/—^* mice were infected with 10^7^ *L. infantum* MEC by IP route, after 2 weeks post infection the animals were euthanized and the livers were used to histological analysis. Livers from non-infected (N.I) and infected C57BL/6 (A, B), *Casp1/11^—/—^* (C), *Il18^—/—^* (D), *Il1b^—/—^* (E) mice were collected after 2 weeks post infection. Granuloma size was quantified in *Casp1/11^—/—^* (F), *Il18^—/—^*(G), *Il1b^—/—^* (H) mice using Image J. Each dot represents a granuloma from a total of 9 (*Il18^—/—^),*10 (*Casp1/11^—/—^*) or 14 (*Il1b^—/—^*) animals. Set of 2 or 3 independent experiments were plotted together. Representative H&E images of granulomas in liver tissues. *, *P* <0.05; **, *P* <0.005; ***, *P* <0.0005; ****, *P* <0.00005.

### NLRP3 inflammasome activation occurs in parasite-infected macrophages and Kupffer cells during hepatic granuloma formation

To further validate inflammasome activation in response to *L. infantum* infection, we performed complementary in vitro experiments using bone marrow-derived macrophages (BMDMs) infected with the NLC strain. Similar to our observations in vivo, infected BMDMs displayed increased NLRP3 puncta formation (**Figure S7A-B**), indicating activation of the NLRP3 inflammasome. In addition, infected macrophages secreted elevated levels of IL-1β following LPS priming, whereas IL-1β production was abolished in *Nlrp3^—/—^* and *Casp1/11^—/—^* cells (**Figure S7C**). These findings further support the ability of *L. infantum* to activate the canonical NLRP3 inflammasome pathway in macrophages.

To directly investigate inflammasome activation in infected cells, we generated tdTomato-expressing *L. infantum* parasites. The tdTomato construct was cloned into the pSSU_neo vector (**Figure S8A**), and parasite transfectants were validated by PCR and fluorescence microscopy (**Figure S8B-C**). Importantly, fluorescent parasites retained their infectivity, as demonstrated by efficient infection of BMDMs in vitro and by their ability to establish infection in the liver and spleen of mice at levels comparable to those observed with wild-type parasites (**Figure S8D-F**). Using these fluorescent parasites, we evaluated intracellular caspase-1 activation by FAM-YVAD-FLICA staining. Infection with *L. infantum* induced robust caspase-1 activation in macrophages (**Figure S9A-B**). Moreover, when infected and non-infected cells were analyzed separately, caspase-1 activation was detected almost exclusively in parasite-containing cells and was dependent on NLRP3 expression (**Figure S9C-E**). Together, these findings provide additional evidence that *L. infantum* directly activates the NLRP3 inflammasome in infected macrophages, supporting our observations in hepatic granulomas during visceral infection.

Since Kupffer cells are thought to serve as a nidus for hepatic granuloma formation during visceral leishmaniasis, we next investigated whether they harbor *L. infantum* parasites in vivo. Using tdTomato-expressing NLC *L. infantum* parasites and intravital microscopy, we observed that parasites were predominantly localized within Kupffer cells and were surrounded by recruited inflammatory cells, forming organized granuloma in the liver (**Figure 4J). Figure S8A-B** show additional examples of tdTomato-expressing *L. infantum* residing within Kupffer cells located in hepatic granulomas. Together with the detection of NLRP3 activation in infected macrophages and hepatic granulomas, these findings support a model in which inflammasome activation occurs within parasite-containing cells and contributes to the organization of the granulomatous response during *L. infantum* infection.

### NLRP3 regulates the cellular composition and organization of hepatic granulomas during *L. infantum* infection

To better understand how NLRP3 contributes to hepatic granuloma organization during *L. infantum* infection, we performed intravital microscopy in infected C57BL/6 and *Nlrp3^—/—^* mice. Prior to imaging, animals were treated with anti-Ly6G, anti-CD45r, and anti-F4/80 antibodies to visualize neutrophils, B cells, and macrophages, respectively. Consistent with our previous observations, hepatic granulomas contained Ly6G+ neutrophils, CD45r+ B cells, and F4/80+ macrophages (**Figure 6A-B, Figure S10A, B**). As expected, *Nlrp3^—/—^* mice developed significantly smaller granulomas than infected C57BL/6 animals (**Figure 6C**). In addition, granulomas from *Nlrp3^—/—^*mice displayed reduced numbers of CD45r+ and Ly6G+ cells (**Figure 6D-F**), whereas the number of F4/80+ cells was not altered. Interestingly, infected *Nlrp3^—/—^* mice exhibited a reduced F4/80+ area within granulomas (**Figure 6G**), suggesting altered activation or expansion of these cells despite their preserved numbers. Consistent with this observation, the total number of F4/80+ cells in the liver was similar between genotypes (**Figure 6H**). In contrast, the total number of CD45r+ cells, but not Ly6G+ cells, was reduced in infected *Nlrp3^—/—^* mice compared with infected C57BL/6 animals (**Figure 6I-J**). Together, these findings indicate that NLRP3 contributes to the proper cellular organization of hepatic granulomas by promoting the accumulation of specific leukocyte populations within these structures.

**Figure 6.**
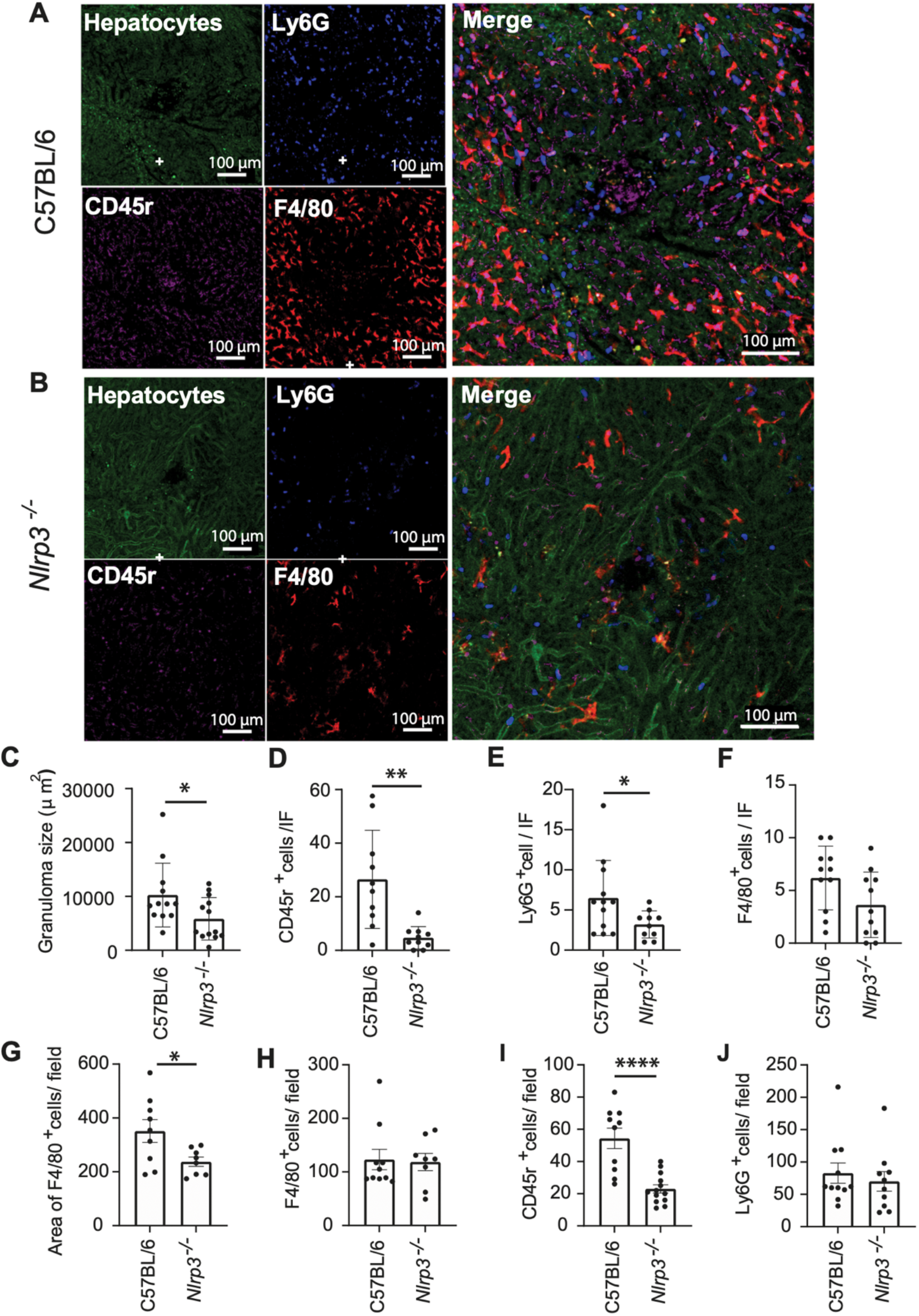
Intravital microscopy reveals NLRP3-dependent granuloma organization and recruitment of neutrophils, B cells, and monocytes during *L. infantum* infection. C57BL/6 and *Nlrp3^—/—^* animals were infected with 10^7^ MEC of *L. infantum* via IP. Between 2 and 3 weeks after infection, the animals were anesthetized and treated with anti-Ly6G, anti-F4/80 and anti-CD45r antibodies. Images of the animals’ livers were obtained using an inverted Nikon Eclipse Ti microscope coupled to an A1R scanning head (Nikon) without alterations. Representative images of the liver of C57BL/6 (A) and *Nlrp3^—/—^* (B) animals infected with *L. infantum*. Granuloma size (C), as well as the number of CD45r+ cells (D), Ly6G+ cells (E) and F4/80+ cells (F) within granulomas were quantified. The area of F4/80+ cells (G) and total number of F4/80+ cells (H), CD45r+ cells (I) and Ly6G+ cells (J) were also quantified. Digital quantification was performed by Volocity (6.3; PerkinElmer) and NIS-Elements (Nikon Instruments). *, *P* <0.05; **, *P* <0.005; ***, *P* <0.0005; ****, *P* <0.00005.

### Lymphocytes, CCR2-dependent monocytes, and neutrophils contribute to hepatic granuloma development during *L. infantum* infection

Since our intravital imaging analyses revealed that hepatic granulomas were composed of multiple leukocyte populations, including monocytes/macrophages, neutrophils, and lymphocytes, we next sought to determine whether these cells contribute to granuloma development during *L. infantum* infection. To address this question, we infected *Rag1^—/—^*mice, which lack mature T and B lymphocytes, and *Ccr2^—/—^* mice, which display impaired monocyte mobilization from the bone marrow to inflamed tissues. Both *Rag1^—/—^* and *Ccr2^—/—^* mice developed significantly smaller granulomas than infected C57BL/6 animals (**Figure 7A-D, G-H**), indicating that lymphocytes and CCR2-dependent monocyte recruitment contribute to granuloma development. Because our previous analyses also identified neutrophils as a prominent component of hepatic granulomas, we next investigated their contribution to this process. Neutrophils were depleted in C57BL/6 mice using the anti-Ly6G antibody (1A8), whereas control animals received the corresponding isotype control antibody (2A3). Neutrophil depletion resulted in a significant reduction in granuloma size following *L. infantum* infection (**Figure 7E-F, I**). The efficiency of neutrophil depletion was confirmed 24 hours after infection, when Ly6G+ cells were virtually absent in animals treated with the 1A8 antibody (**Figure S11A-B**). Together, these findings demonstrate that lymphocytes, CCR2-dependent monocytes, and neutrophils all contribute to the establishment and development of hepatic granulomas during *L. infantum* infection.

**Figure 7.**
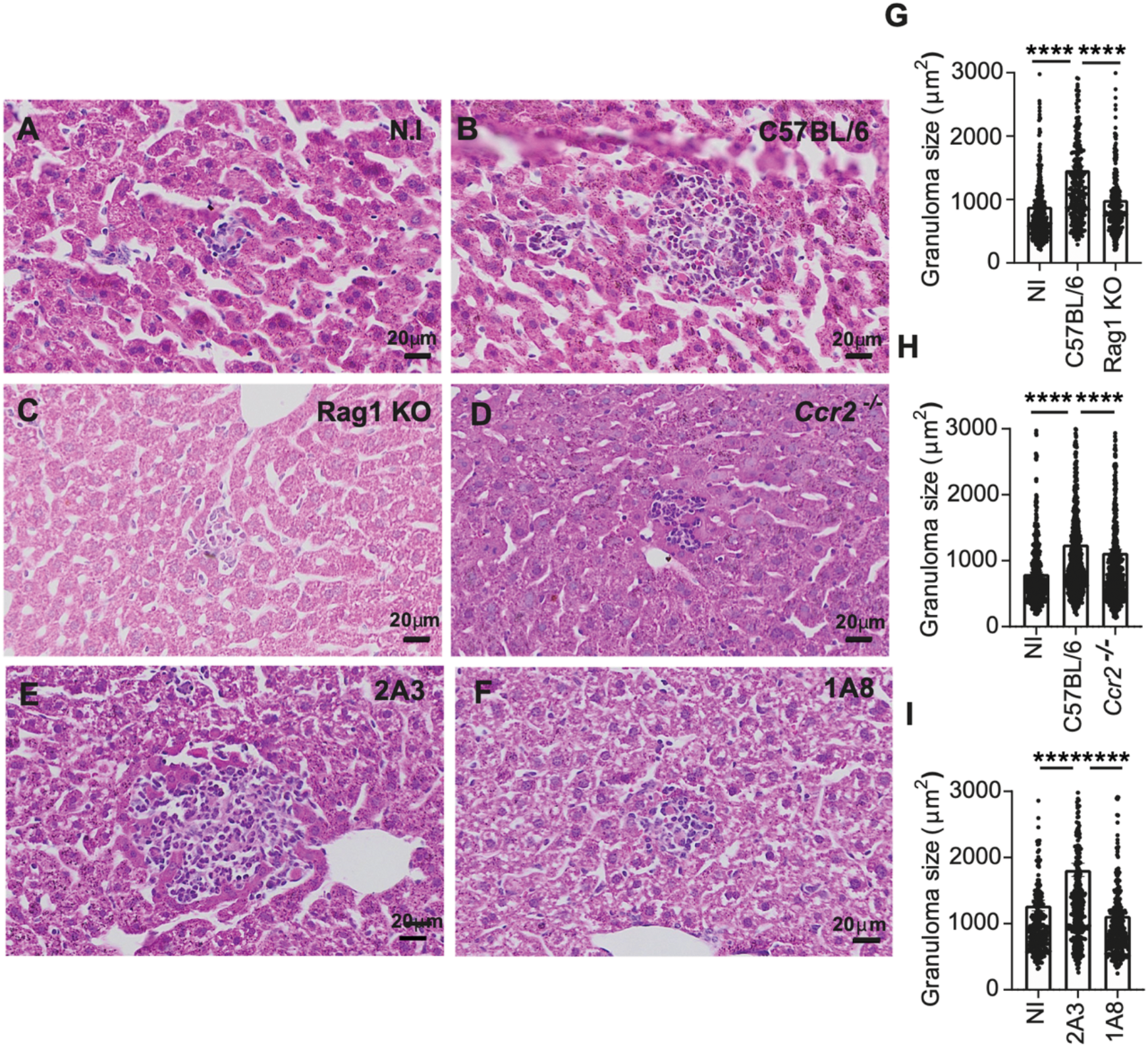
Lymphocytes, CCR2-dependent monocytes, and neutrophils promote hepatic granuloma development during *L. infantum* infection. C57BL/6, Rag1 KO and *Ccr2^—/—^*, were infected with 10^7^ MEC of *L. infantum* NLC strain by IP route, after 2 weeks post infection the animals were euthanized and the livers were used to histological analysis. Livers from non-infected (N.I) and infected C57BL/6 (A, B), Rag1 KO (C), *Ccr2^—/—^* (D) mice were collected. Granuloma size was quantified in Rag1 KO (G) and *Ccr2^—/—^*(H) mice using Image J. Each dot represents a granuloma from a total of 10 (*Ccr2^—/—^*) or 12 (Rag1 KO) animals. Set of 2 independent experiments were plotted together. C57BL/6 mice were infected with 10^7^ *L. infantum* MEC via IP, after 1 week of infection the animals were treated with the rat IgG2a isotype control (2A3) or anti-mouse Ly6G (1A8) antibody. Two weeks after infection, the animals were sacrificed and the livers were used for histological analysis. Representative H&E images of granulomas in the livers of infected mice treated with 2A3 antibody (E) and 1A8 antibody (F). Granuloma size was quantified (I) using Image J. Each dot represents a granuloma from a total of 10 animals per group. Set of 2 independent experiments were plotted together. *, *P* <0.05; **, *P* <0.005; ***, *P* <0.0005; ****, *P* <0.00005.

### NLRP3-dependent granuloma organization is associated with host resistance to *L. infantum* infection

Given that NLRP3 deficiency impaired hepatic granuloma development and altered the recruitment of immune cells into these structures, we next investigated whether NLRP3-dependent granuloma organization contributes to host resistance against *L. infantum* infection. To address this question, *Nlrp3^+/+^*, *Nlrp3^+/—^*, and *Nlrp3^—/—^* littermate mice were infected intraperitoneally with 10^7^ *L. infantum* parasites, and parasite burdens were quantified in the liver and spleen. *Nlrp3^—/—^*mice exhibited significantly higher parasite loads in both organs at weeks 3 and 4 post-infection compared with *Nlrp3^+/+^* and *Nlrp3^+/—^*animals (**Figure 8A-D**), indicating increased susceptibility to infection in the absence of NLRP3. To further determine whether this phenotype was associated with inflammasome signaling, we evaluated parasite burdens in *Nlrp3^—/—^*, *Casp1^—/—^,* and *Casp1/11^—/—^* mice. Consistent with the results observed in *Nlrp3^—/—^* animals, mice deficient in inflammatory caspases displayed significantly increased hepatic parasite burdens at both 3 and 4 weeks post-infection compared with C57BL/6 controls (**Figure 8E-F**). Similarly, *Nlrp3^—/—^*, *Casp1^—/—^,* and *Casp1/11^—/—^* mice exhibited increased splenic parasite burdens at week 3 post-infection, although these differences were no longer evident at week 4 (**Figure 8G-H**). No differences in parasite loads were detected at week 2 post-infection between *Nlrp3^—/—^* and C57BL/6 mice (**Figure S12**), suggesting that the effects of NLRP3 deficiency become evident during the phase of granuloma expansion and maturation.

**Figure 8.**
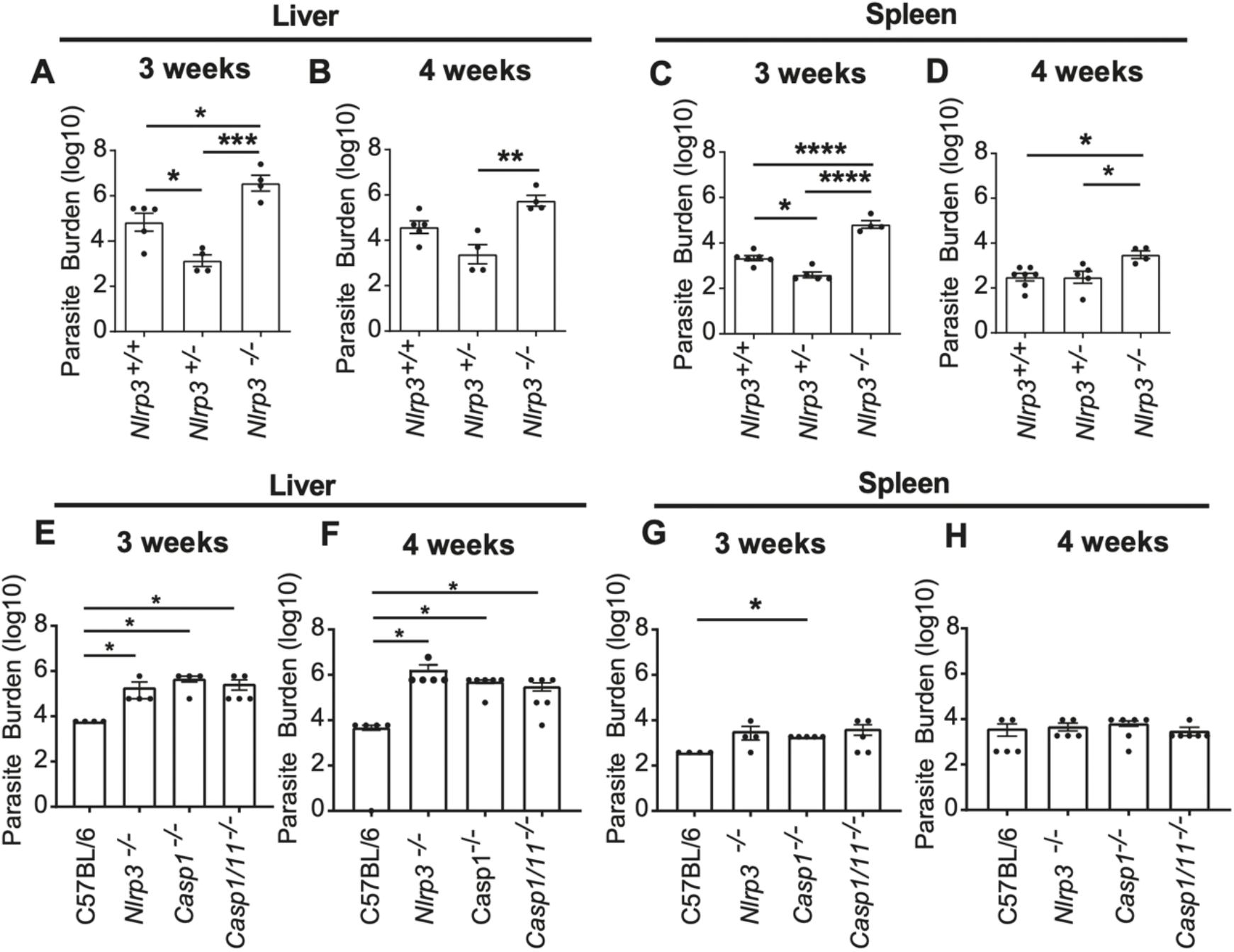
NLRP3 and inflammatory caspases contribute to control of *L. infantum* in the liver and spleen. *Nlrp3*^+/+^, *Nlrp3^+/—^* and *Nlrp3^—/—^* from litter mate control (*Nlrp3*^+/—^ x *Nlrp3*^+/—^) were genotyped and infected with 10^7^ MEC of *L. infantum* NLC strain. Livers (A, B) and spleens (C, D) were collected after 3 weeks and 4 weeks post infection and used to measure parasite burden by limited dilution assay. *Nlrp3^—/—^*, *Casp1^—/—^*, *Casp1/11^—/—^* mice were infected with 10^7^ *L. infantum* MEC by IP route. 3- and 4-weeks post infection, the animals were euthanized and the livers (E, F) and spleen (G, H) were used to measure parasite burden by limited dilution assay. Quantification of the parasitic load was carried out by a limiting dilution assay. *, *P* <0.05; **, *P* <0.005; ***, *P* <0.0005; ****, *P* <0.00005.

To further investigate NLRP3-mediated resistance to infection, we monitored parasite replication using luciferase-expressing NLC *L. infantum* parasites and the IVIS Spectrum Optical Imaging Platform. Longitudinal imaging revealed increased luminescence in *Nlrp3^—/—^* mice at 3 weeks post-infection (w.p.i.), whereas no differences were observed at 1 or 2 w.p.i. (**Figure 9A-E**). Luciferase activity was also measured in liver samples at a 10^-5^ dilution at 3 w.p.i. Consistent with the in vivo imaging, luciferase activity was increased in the livers of *Nlrp3^—/—^*mice compared with C57BL/6 mice (**Figure 9F**). Limiting dilution assay of the same experiment further confirmed an increased parasite burden in the livers of *Nlrp3^—/—^* mice (**Figure 9G**). In this experiment we also quantified parasite load by qPCR and revealed an increased number of *Leishmania* parasites per cell in the livers of *Nlrp3^—/—^*mice (**Figure 9H**). Together, these complementary approaches confirm that NLRP3 contributes to the control of *L. infantum* replication, with differences becoming evident at 3 weeks after infection, coinciding with the period of granuloma development.

**Figure 9.**
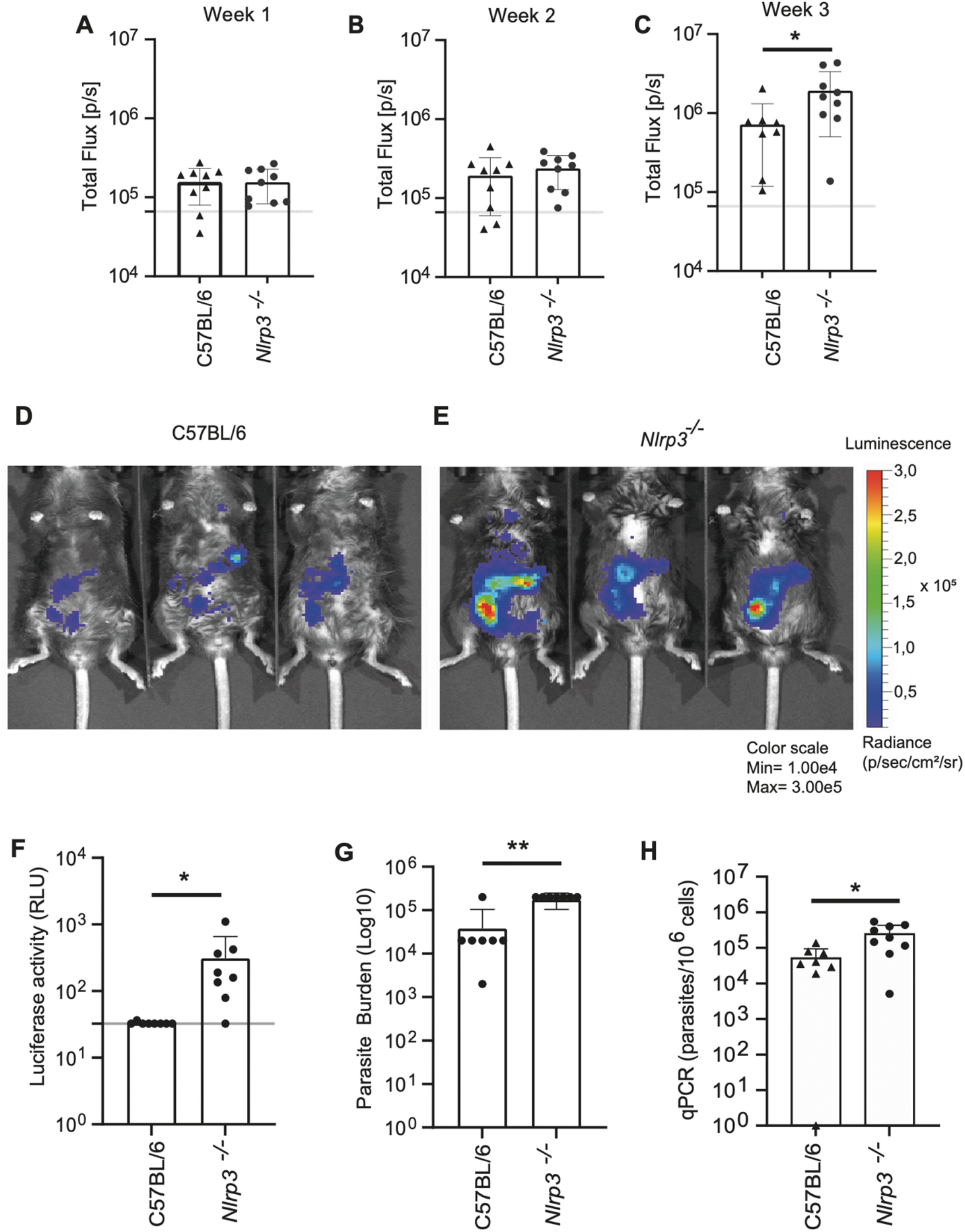
NLRP3 promotes hepatic control of *L. infantum* infection assessed by bioluminescence imaging, limiting dilution, and qPCR. C57BL/6 and *Nlrp3^—/—^* mice were infected with 10^7^ MEC luciferase-expressing NLC *L. infantum* parasites. After 1 (A), 2 (B) and 3 (C) weeks post infection animals were observed in IVIS Spectrum Optical Imaging Platform. Representative imaging of IVIS Spectrum Optical Imaging Platform images of C57BL/6 and *Nlrp3^—/—^* mice (D, E). After 3 weeks post infection, the animals were euthanized and the liver was collected. Luciferase activity was measured using a 10^-5^ dilution of liver homogenate. The gray line represents the cutoff used, established using a non-infected liver (F). Liver homogenate was also used for limited dilution assay to determine parasite burden (G). Parasite burden in liver was also determined by qPCR (H). *, *P* <0.05; **, *P* <0.005; ***, *P* <0.0005; ****, *P* <0.00005.

### NLRP3-dependent granulomatous inflammation is associated with both hepatic pathology and host protection during *L. infantum* infection

Since NLRP3 inflammasome signaling promoted granuloma organization and host resistance, we next investigated how these effects influenced hepatic pathology. This question is particularly relevant because previous studies have attributed both protective and detrimental functions to inflammasome activation during cutaneous leishmaniasis (reviewed by (Harrington and Gurung, 2020; Zamboni and Sacks, 2019). Therefore, we performed a detailed histopathological analysis to determine whether NLRP3-dependent granulomatous inflammation was associated with increased liver injury or, alternatively, represented a protective and organized tissue response.

Histopathological analyses were performed by a board-certified pathologist in liver sections obtained from infected and non-infected C57BL/6 and *Nlrp3^—/—^* mice. As expected, livers from non-infected animals of both genotypes displayed preserved hepatic architecture and no significant histopathological alterations (**Figure 10A-B**). Following infection, both C57BL/6 and *Nlrp3^—/—^* mice developed mild hepatic changes, including cytoplasmic vacuolization, accumulation of lipid droplets consistent with microvesicular steatosis, and increased frequencies of binucleated hepatocytes and polyploid nuclei (**Figure 10C-F**). However, the severity of these alterations was generally lower in *Nlrp3^—/—^* mice, particularly at 3 weeks post-infection, when these animals exhibited the lowest overall histopathological scores (**Figure 10H**). In addition to mild parenchymal alterations, infected animals developed portal inflammatory infiltrates and lobular granulomatous lesions. Notably, granulomatous inflammation was less prominent in *Nlrp3^—/—^* mice, particularly at 2 weeks post-infection (**Figure 10G**). Consistent with our previous quantitative analyses, granulomas from *Nlrp3^—/—^*mice were smaller and exhibited a looser cellular organization compared with those observed in infected C57BL/6 mice (**Figure 10G, I-L**). The histopathological criteria used for lesion scoring are detailed in the supplementary material (**Figure S13**).

**Figure 10.**
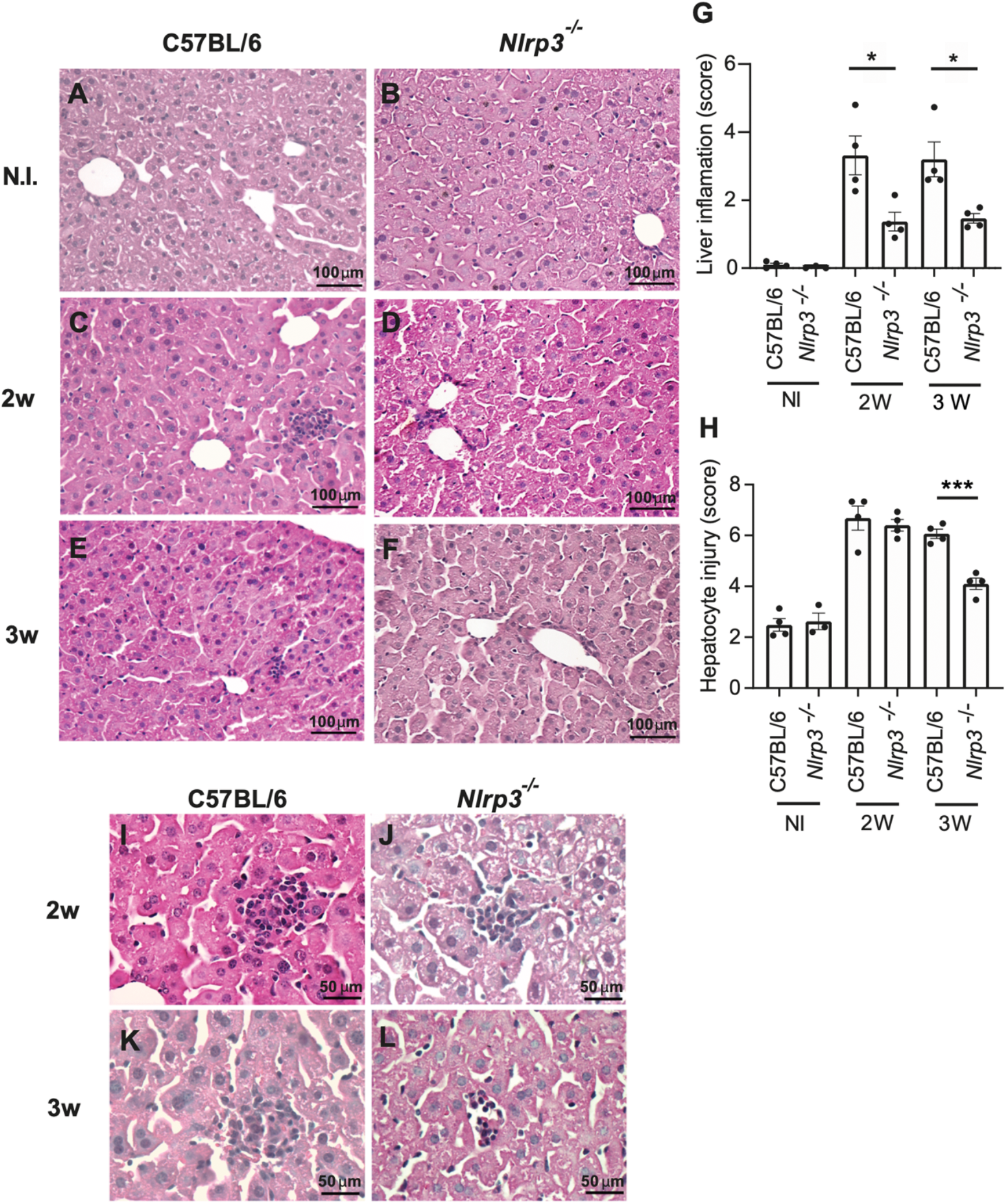
NLRP3 promotes hepatic inflammation and granulomatous pathology during *L. infantum* infection. Representative images of the histological sections of liver from C57BL/6 and *Nlrp3^—/—^* infected or non-infected mice. Liver from non-infected C57BL/6 and *Nlrp3^—/—^*mouse (A, B); Liver from 2 weeks-infected C57BL/6 and *Nlrp3^—/—^* mouse (C, D); Liver from 3 weeks-infected C57BL/6 and *Nlrp3^—/—^* mouse (E, F). Liver inflammation score (G). Degree of hepatocyte injury (H). Representative images of the histological sections of liver granuloma from C57BL/6 and *Nlrp3^—/—^*infected mice after 2 weeks (I, J) and 3 weeks (K, L) of infection. HE; Hematoxylin-eosine staining. *, *P* <0.05; **, *P* <0.005; ***, *P* <0.0005.

Together, these findings indicate that NLRP3 deficiency reduces hepatic inflammation and tissue alterations during *L. infantum* infection despite increased parasite burdens. While inflammasome activation has been associated with both protective and pathogenic outcomes in different models of leishmaniasis, our results demonstrate that, in this model of resistance, NLRP3-dependent inflammation is linked to improved parasite control and the formation of organized hepatic granulomas. Therefore, although NLRP3 signaling contributes to local inflammatory pathology, these responses appear to represent a beneficial trade-off that favors host protection.

## Discussion

The mechanisms that determine whether granulomatous inflammation develops into an organized and protective immune structure remain incompletely understood. Here, we identify the NLRP3 inflammasome as an important regulator of hepatic granuloma maturation and host resistance during visceral leishmaniasis. Although NLRP3 was dispensable for the initial establishment of granulomatous lesions, its absence impaired granuloma expansion and cellular organization and was consistently associated with defective control of *L. infantum*. The latter phenotype was confirmed using complementary approaches, including longitudinal bioluminescence imaging, limiting dilution, and qPCR, and became evident during the period of active granuloma development. Together, these findings support a model in which NLRP3 contributes to protective immunity not simply by amplifying inflammation, but by promoting the development of an organized granulomatous response capable of supporting effective parasite control.

An important finding of our study is the evidence of systemic inflammasome activation in patients with active visceral leishmaniasis. Although increased circulating IL-1β has been reported in VL (Araujo-Santos et al., 2017; Babaloo et al., 2020), IL-1β alone is not a definitive indicator of inflammasome activation because it can also be processed through inflammasome-independent mechanisms, including neutrophil-derived proteases (Coeshott et al., 1999; Guma et al., 2009; Joosten et al., 2009). In our cohort, the concomitant increase in IL-18 and cleaved caspase-1 (Casp1 p20), together with elevated IL-1β, provides stronger evidence that inflammasome activation occurs during active human disease. This observation extends previous studies linking IL-18 polymorphisms, peripheral blood cell responses, and experimental infection to inflammasome-associated pathways during *Leishmania* infection (Ahmadpour et al., 2016; Haeberlein et al., 2010; Kumar et al., 2014; Lima-Junior et al., 2013; Moravej et al., 2013; Santos et al., 2018), and establishes a clinical context for the mechanistic role of NLRP3 identified in our experimental model.

Previous studies have established that *Leishmania* can engage inflammasome pathways in macrophages, although their consequences vary with parasite species and infection context (Charmoy et al., 2016; Chaves et al., 2019; de Carvalho et al., 2019; de Sa et al., 2023; Gupta et al., 2017; Gurung et al., 2015; Lima-Junior et al., 2013). In visceral leishmaniasis’, this response occurs within a highly dynamic macrophage compartment’, including resident Kupffer cells seed hepatic granulomas’, whereas recruited monocytes progressively accumulate and differentiate as these structures mature (Pessenda et al., 2025). Our transcriptomic analyses place inflammasome signaling within this cellular transition. Inflammasome-associated transcripts were enriched in hepatic macrophages and spatially concentrated in granuloma-associated regions (Dey et al., 2026), while single-cell analysis of an independent *L. infantum* dataset revealed the strongest signature in recruited and transitioning monocyte-derived populations rather than resident Kupffer cell states (Pessenda et al., 2025). Thus, inflammasome-associated gene expression appears linked to specific macrophage states acquired during granuloma maturation rather than uniformly distributed across hepatic macrophages. Although transcriptomic signatures do not demonstrate biochemical inflammasome activation, their spatial and cellular distribution, together with our experimental evidence of NLRP3 activation, supports a model in which inflammasome signaling is integrated into the macrophage remodeling that accompanies granuloma development.

Previous studies have linked inflammasome-associated pathways to granulomatous diseases and mycobacterial infection, but these studies have largely focused on inflammation, cytokine production, fibrosis, or tissue pathology rather than on the mechanisms that determine granuloma architecture (Denis, 1994; Huppertz et al., 2020; Juffermans et al., 2000; Master et al., 2008; Sugawara et al., 1999; Theobald et al., 2024). Our findings extend this framework by identifying the NLRP3-Caspase-1/11-IL-18 axis as a regulator of granuloma maturation and cellular organization. Notably, preserved granuloma development in *Il1b^—/—^* mice indicates that this function preferentially involves IL-18 rather than reflecting a general consequence of IL-1 family cytokine production. This is consistent and contributes to mechanistically explain previous studies showing that IL-18 contributes to resistance during experimental *L. donovani* infection and can promote IFN-γ-dependent antimicrobial responses (Mullen et al., 2006; Murray et al., 2006; Vieira et al., 2024). In parallel, the requirement for lymphocytes, CCR2-dependent monocytes, and neutrophils for optimal granuloma development, together with their altered accumulation in *Nlrp3^—/—^* lesions, suggests that NLRP3 functions upstream of cellular events that shape the granulomatous niche. Whether IL-18 directly controls leukocyte recruitment or instead acts through activation and cross-talk among granuloma-associated cells remains to be determined.

A longstanding concept in experimental visceral leishmaniasis is that hepatic granuloma maturation is closely linked to the acquisition of parasite control (McElrath et al., 1988; Murray, 2001). Our findings place NLRP3 within this protective axis. *Nlrp3^—/—^* mice developed smaller and less organized granulomas and exhibited impaired hepatic parasite control, a phenotype independently confirmed by longitudinal bioluminescence imaging, limiting dilution, and qPCR. Importantly, differences in parasite burden became evident at 3 weeks after infection, coinciding with the period of active granuloma maturation. This temporal association supports the possibility that NLRP3-dependent granuloma organization contributes to efficient local parasite control. However, our data do not establish that defective granuloma architecture is itself the cause of increased parasite burden, and NLRP3 may additionally influence antimicrobial functions or immune-cell interactions independently of granuloma organization. Thus, NLRP3 emerges as a link between inflammatory sensing, granuloma maturation, and protective immunity during visceral leishmaniasis.

The protective function of NLRP3 was accompanied by increased hepatic inflammation, highlighting the context-dependent consequences of inflammasome signaling. *Nlrp3^—/—^* mice exhibited reduced inflammatory and histopathological alterations despite impaired parasite control, indicating that the inflammatory response promoted by NLRP3 is not simply detrimental in this setting. This duality is consistent with the broader concept that inflammasome activation can contribute to either host defense or tissue injury depending on the infectious and tissue context (Broz and Dixit, 2016; Guo et al., 2015; Swanson et al., 2019). In visceral leishmaniasis, our findings suggest that part of the inflammatory cost associated with NLRP3 activation accompanies the formation of a more organized and protective granulomatous response. Overall, our study identifies granuloma organization as a previously underappreciated effector function of inflammasome signaling and supports a model in which NLRP3, acting through Caspase-1/11 and IL-18 and in concert with recruited monocytes, neutrophils, and lymphocytes, contributes to protective immunity by shaping the cellular and spatial architecture of granulomatous responses within tissues.

## Materials and Methods

### Participants

Serum samples from healthy individuals and VL patients were kindly provided by Dr. Carlos Henrique Nery Costa, from the Federal University of Piauí. At the Natan Portela Institute of Tropical Diseases (IDTNP), a reference hospital for infectious diseases, 100 individuals diagnosed with visceral leishmaniasis were treated. Participants were selected based on diagnostic criteria, specifically those with a confirmed diagnosis of VL by culture, with parasites cryopreserved in liquid nitrogen. Clinical and laboratory data were carefully collected from patients’ medical records. For each isolate, a 250 μL serum aliquot was collected prior to treatment initiation and stored at −20 °C to preserve its immunological integrity for subsequent analyses. The protocol, together with informed consent obtained from all participants or their legal guardians, was initially approved by the Research Ethics Committee of the Federal University of Piauí under number 0116/2005. Although initial ethical approval was granted in 2005, the project has since been renewed and continuously monitored to ensure compliance with current ethical standards. The study was conducted in accordance with the principles established in the Declaration of Helsinki, governing research involving human subjects.

### Quantification of cytokines and caspase-1 in human serum

Serum cytokine levels in VL patients and control subjects were quantified using the BD™ CBA Human Th1/Th2/Th17 Cytokine Kit (Cat. No. 560484) and the BD™ CBA Human Inflammatory Cytokine Kit (Cat. No. 551811) (BD Biosciences), following the manufacturer’s instructions. Samples were acquired on a BD Accuri C6 flow cytometer, and data were analyzed using FlowJo software (Tree Star, Ashland, OR, USA). Quantification of GSDMD, caspase-1 p20, and IL-18 was performed by ELISA, using the Abcam ab272463 (GSDMD), R&D Systems DCA100 (Caspase-1 p20), and R&D Systems DY318 (IL-18) kits. Samples were read on a microplate reader, and concentrations were determined from standard curves generated with specific recombinant standards. ELISA data were analyzed using SoftMax Pro software (Molecular Devices, San Jose, CA, USA).

### Single-cell RNA-seq analysis

Single-cell RNA-seq data were obtained from Pessenda et al. (Pessenda et al., 2025) for *Leishmania infantum* data and from Dey et al. (Dey et al., 2026) for *Leishmania donovani* data. We used the processed Seurat object and cell annotations publicly provided by the authors, preserving the original UMAP embedding and annotation fields. Per-cell inflammasome signature activity was quantified using AUCell (Andreatta and Carmona, 2021) based on the genes Nlrp3, Pycard, Casp1, Il1b, and Il18. Downstream analyses and visualizations were performed in R using Seurat, AUCell, dplyr, and ggplot2.

### Spatial transcriptomics analysis

Visium spatial transcriptomics data were obtained from Dey et al. (Dey et al., 2026). We analyzed the infected Visium sections X334.I2, X334.I4, X345.I3, and X345.I5, preserving the spatial coordinates and the transcriptomic clustering provided by the original study. Cluster 4 was used as the main granuloma-associated region of interest, based on the original study showing that RNA_4 overlapped histologically defined granulomas. Inflammasome activity was scored at the spot level using AUCell with *Nlrp3*, *Pycard*, *Casp1*, *Il1b*, and *Il18*. For statistical testing, we compared cluster 4 against all other spots within each infected section by calculating, for each sample, the difference between the mean inflammasome AUCell score in cluster 4 and the mean score in non-cluster-4 spots. Significance was assessed by a within-sample permutation test with 10,000 permutations, in which cluster 4 labels were randomly shuffled within each section while preserving the number of cluster 4 spots per sample. The one-sided enrichment p value was calculated as the fraction of permuted statistics greater than or equal to the observed statistic.

### *Leishmania* strains and culture

*Leishmania infantum* strain IOCL 3241 (MHOM/BR/2005/NLC), Luciferase-expressing *Leishmania infantum* IOCL 3241, *Leishmania infantum* strain Pp75 (MHOM/BR/1974/PP75) and *Leishmania donovani* strain LV9 (MHOM/IN/80/DD8) were used in this work. *Leishmania spp* were grown in Schineider supplemented with 10% fetal bovine serum (Gibco), 100 U/mL penicillin/streptomycin (Sigma) and 0.025% biopterine (Sigma), or 199 Media (Gibco), supplemented with 10% heat-inactivated fetal bovine serum (FBS), 40 mM HEPES, 0.1 mM adenine, 5 mg/L hemin, 1 mg/L biotin, 70 U/mL penicillin and 70 µg/mL streptomycin. The parasites were routinely grown in sterile culture bottles and kept at 27 °C in an incubator.

### Preparation of *Leishmania spp* for infection

For in vitro infection, the stationary phase culture of parasites was centrifuged at 130 x g for 3 minutes to eliminate the clumped parasites. The supernatant was centrifuged at 3200 x g for 10 minutes to concentrate procyclic and metacyclic promastigotes in the pellet. The pellet was resuspended and used for infection. For in vivo experiments, we generated a Metacyclic-enriched culture (MEC) for infection, which contains lower frequency of procycliclic promastigotes in the final parasite population. For this, a first centrifugation of 130 x g was performed for 3 minutes. The supernatant was used for a second centrifugation of 1250 x g for 5 minutes to precipitate (and eliminate) part of the procyclic promastigotes. Finally, a final centrifugation of 3200 x g for 10 minutes was preformed to concentrate metacyclic promastigotes in the pellet, that was further resuspended to obtain the metacyclic enhanced culture.

### Generation of fluorescent *L. infantum* parasites in the NLC strain

To obtain fluorescent parasites, we carried out the construction of *L. infantum* NLC dTomato strain. For that, the plasmid pSSUneoTdTOMATO (Beattie et al., 2010) was linearized with the enzymes PacI and PmeI (New England Biolabs). About 5 µg of linearized DNA were used to transfect *L. infantum* NLC. According sequence homology, the construction was directed to the locus that encodes the smaller subunit of the ribosomal RNA (18S), under control of RNA polymerase I promoter (constitutive transcription). The products obtained were selected in a solid 199 medium (Sigma), containing 30 µg/ml of G418. The integration of the construction in the SSU locus was confirmed by polymerase chain reaction (PCR), whose strategy was to use a primer that binds only at the construction and another only at the locus, in order to obtain amplification only when the construction was integrated in the locus of interest. TdTOMATO expression was also verified by flow cytometry and fluorescence microscopy.

### Mice

All animal experiments were performed in accordance with the guidelines of the Ethics Committee on the Use of Animals (CEUA) and approved under the protocol number (1147/2022). C57BL/6J mice were obtained from Jackson (Jax 000664), breed and maintained in Central Animal Facility of the School of Medicine of Ribeirão Preto (FMRP), University of São Paulo (USP). NLRP3-deficient mice (*Nlrp3^-/-^*) were also from Jackson (JAX 017969). *Nlrp3^-/-^* and C57BL/6 were breed to generate heterozygous F1 mice that were intercrossed to generate F2 mouse population, containing *Nlrp3^-/-^* and *Nlrp3^+/+^*littermate controls. *Casp1/11^-/-^* mice (Kuida et al., 1995) were backcrossed with C57BL/6 for eight generations. *Casp1^-/-^* mice were generated in the C57BL/6 background as described (Rauch et al., 2017). *Rag1^—/—^* (JAX 002216) and *Ccr2^—/—^* (JAX 004999), *Il1r1^—/—^* (JAX 003245) and *Il18^—/—^* (JAX 004130) mice were obtained from Jackson backcrossed with C57BL/6J and maintained at the institutional facilities. Mice of both sexes, between 6 and 9 weeks of age, were maintained in institutional animal facility in microisolators, with light-dark cycle every 12 hours, temperature maintained between 22 and 25°C and received water and feed ad libitum. Adult male and female C57BL/6 Ccr2RFP Cx3cr1GFP mice (6 weeks old) were taken from the Bioterism Center of the Federal University of Minas Gerais (Brazil). Animals were raised by breeders with 2 female mice per male per cage. All animals were housed in acrylic cages with a filtered air system (Alesco; 5 mice/cage) in a specific pathogen-free conventional facility at the Federal University of Minas Gerais with water and food provided ad libitum and light-dark cycle (12 /12h). All experiments with mice were approved by the Animal Ethics Committee of the Federal University of Minas Gerais (registration number 034/2017), following international guidelines for animal care.

### Culture and infection of BMDMs

C57BL/6, *Nlrp3^-/-^*, *Casp1^-/-^* and *Casp1/11^-/-^* mice were euthanized and femurs and tibias carefully removed. The bones were washed with 70% alcohol and maintained in Phosphate Buffered Saline (PBS) (Gibco). Bones were sectioned and their inner marrow was removed with the aid of a syringe with a needle and RPMI medium (Gibco). The freshly harvested cells were diluted in 10 mL of RPMI medium enriched with 20 % fetal bovine serum (Gibco), 100 U/mL penicillin/streptomycin (brand) and 30 % supernatant from L929 cells in untreated Petri dishes (100 mm x 20 mm) and then incubated in an incubator at 37°C injected with 5% CO_2_. The cells received an additional 10 mL of R20/30 after 4 days. On the 7th day, bone marrow-derived macrophages (BMDMs) were removed using 5 mL of pre-heated (37°C) sterile PBS to wash and remove only non-adherent cells. Then, 10 mL of cold sterile PBS was added. The plates were taken to the refrigerator for 10 minutes, then kept at room temperature for another 5 minutes. The cells were released by washing with the aid of a pipettor, then the total volume was centrifuged at 200 x g for 5 minutes. The macrophage pellet was resuspended in RPMI medium supplemented with 10% fetal bovine serum and 2 mM L-glutamine and counted in a Neubauer chamber. Then, 5 x 10^4^ BMDMs were plated on 96-well plates/well or 2 x 10^5^ cells on 24-well plates/well with round millimeter coverslips at the bottom. Then these macrophages were infected with *Leishmania infantum* with MOI 5 or MOI 10 for 4 hours in an incubator at 37°C and 5% CO_2_ and so washed twice with PBS. The non and infected cells were maintained in the incubator at 37°C and 5% CO_2._

### Caspase-1 activity measurement

A total of 10^6^ *L. infantum* NLC-infected or not BMDMs were previously plated and then were released with the help of a squeegee. Next, the cells were centrifuged at 200 x g for 5 minutes and then labeled for 30 min with the FLICA carboxyfluorescein reagent (FAM–YVAD–FMK; Immunochemistry Technologies) at 37°C, as recommended by the manufacturer. After, the cells were washed three times with PBS. Acquisition was performed in a flow cytometer (BD Accuri C6; BD Biosciences) and analyzed using FlowJo (Tree Star) software.

### Immunofluorescence staining of infected BMDMs

For staining *L. infantum*-infected or not BMDMs, a total of 5 × 10^5^ BMDMs were plated in 24-well plate with containing round coverslips for 24 h. For fixation of the samples, tissue culture supernatants were removed, and cells were fixed with 4% paraformaldehyde for 20 min at room temperature. Paraformaldehyde was removed, cells were washed with PBS, and then the coverslips were processed for immunofluorescence. For that, cells were blocked and permeabilized using PBS with goat serum and 0.05% saponin for 1 h at room temperature. Next, primary antibodies used were rabbit mAb anti-human NLRP3 (clone D2P5E, batch 2; 1:1,000; Cell Signaling) or rabbit polyclonal antibody anti-human ASC (1:2,000; Adipogen AL177). Antibodies were diluted in blocking solution. After 1 h of incubation, the samples were washed twice with PBS, and secondary antibodies were added and incubated for 1 h at room temperature. Secondary antibodies used were goat anti-rabbit 488 (1:3000; Invitrogen) and goat anti-rabbit 594 (1:3000; Life Technologies). Slides were washed twice with PBS and then mounted using DAPI (1 mM) and ProLong (Invitrogen).

### Mice infection and quantification of parasitic load by limiting dilution assay

The purified and counted parasites were resuspended in sterile PBS (Gibco) for infection of the animals. 10^6^ or 10^7^ parasites in 100 µL of PBS were injected intraperitoneally. At the indicated times after infection, the mice were euthanized to collect the spleen and liver. After being collected, the organs were washed in 70% alcohol and then in sterile PBS, and transferred to 6-well plates containing 5 mL of Schineider medium. With the aid of the plunger of a 5 mL syringe, they were macerated to release of intracellular amastigotes and immune cells. Then, the macerate was transferred to a falcon containing a Cell Strainer of 70 nM (Falcon), the wells were washed with another 10 mL of Shineider medium, which went on to the centrifugation steps. The first centrifugation was 130 x g for 3 minutes, to separate red blood cells and remaining tissues. The supernatant was again centrifuged at 3200 x g for 10 minutes to collect the amastigotes. The pellet was resuspended in a complete Schineider medium to perform the LDA. The plates were incubated for 10 days until the amastigotes were differentiated into promastigotes to be analyzed under the microscope, thus allowing the quantification of the parasitic load.

### *In vivo* bioluminescence imaging

Mice infected with Luciferase-expressing *Leishmania infantum* was monitored by in vivo bioluminescence imaging on days 7, 14 and 21 post-infection. On day 21, mice were euthanized, and spleens and livers were collected for parasite burden determination by LDA or q PCR. *In vivo* bioluminescence imaging was performed using an IVIS Spectrum imaging system (PerkinElmer). At each imaging time point, mice received an intraperitoneal injection of D-luciferin (VivoGlo™, Promega; 100 mg/kg body weight) and were anesthetized with isoflurane (3% for induction and 1% for maintenance). Images were acquired 10 min after D-luciferin administration using an exposure time of 10s. Total photon flux within defined regions of interest (ROIs) was quantified using Living Image software (PerkinElmer) and expressed as photons/s. An uninfected mouse was imaged under the same conditions and served as a negative control for background bioluminescence.

### Luciferase viability assay

A limiting dilution assay was conducted using liver homogenates. Samples were serially diluted (1:10) in M199 medium supplemented as described above in 96-well plates, in duplicate, and incubated at 26 °C for 14 days. Following incubation, aliquots from each well were transferred to 96-well, white, flat-bottom Nunc plates (Thermo Fisher Scientific). Homogenates from an uninfected mouse were processed in parallel and used as negative controls for background luminescence. ONE-Glo™ Luciferase Assay System (Promega) was added according to the manufacturer’s protocol, and the luminescence signal was read using a Varioskan™ LUX Multimode Microplate Reader (Thermo Fisher Scientific). The highest dilution yielding a luminescence signal above background was recorded as the endpoint dilution. Samples with luminescence above background were considered positive, while samples with no detectable signal above background were classified as not detected (ND).

### DNA extraction and quantitative PCR

Genomic DNA was extracted from liver samples using the DNeasy Blood & Tissue Kit (QIAGEN), according to the manufacturer’s instructions. Parasite burden was also determined by quantitative real-time PCR (qPCR) targeting *Leishmania infantum* kinetoplast DNA (kDNA). Reactions were performed in a final volume of 20 µL containing 200 ng of genomic DNA, 0.3 µM of each primer and 0.2 µM of the hydrolysis probe, using TaqMan™ Universal PCR Master Mix (Thermo Fisher Scientific). The following oligonucleotides were used: kDNA_124-145_Fw (5′-CCACCCGGCCCTATTTTATTTTACACC-3′), kDNA_1-23_Rv (5′-CCAAACTTTTCTGGTCTCTCCGGG-3′), and kDNA_P_23-56 (5′-FAM-TAGGGGCGTTCTGCGAAAATCGAAAAATGGGTGC-BHQ1-3′). Thermal cycling consisted of an initial activation step at 95 °C for 5 min, followed by 45 cycles of denaturation at 95 °C for 5 s and annealing/extension at 60 °C for 15 s. A six-point standard curve corresponding to 0.1–10,000 parasites was used for parasite quantification. Host-cell equivalents were determined by qPCR targeting the mouse β-actin (Actb) gene. Reactions were performed in a final volume of 15 µL containing 200 ng of genomic DNA and 0.13 µM of each primer using Power SYBR™ Green PCR Master Mix (Thermo Fisher Scientific). The primers used were Actb-Fw (5′-CGATGCCCTGAGGCTCTTT-3′) and Actb-Rev (5′-TGGATGCCACAGGATTCCAT-3′). Thermal cycling consisted of an initial activation step at 95 °C for 10 min, followed by 45 cycles of denaturation at 95 °C for 15 s and annealing/extension at 60 °C for 15 s. A standard curve generated from serially diluted mouse splenocytes, ranging from 1 to 10,000 cells, was used to determine host-cell equivalents. All reactions were performed on a QuantStudio™ 3 Real-Time PCR System (Applied Biosystems), and all samples and negative controls were analyzed in technical duplicate. Parasite burden was calculated from the kDNA standard curve and normalized to the corresponding number of mouse cells determined by Actb qPCR. Results were expressed as the number of *Leishmania* cells per 10^6^ mouse cells.

### Neutrophils depletion

To determine the importance of neutrophils in *L. infantum* NLC infection by IP route, the animals were depleted or not using 1 mg of anti-Ly6G (1A8, Bioxcell) and isotype control (2A3, Bioxcell) 24 hours prior infection or 1 week after infection. Twenty-four and forty-eight hours post infection, the number of neutrophils at peritoneum were estimated.

### Immunoflorescence and imaging of infected livers

Infected and non-infected mice were perfused with PBS and the livers collected. Then the livers were cut, and fixed in 10% buffered formalin. The tissue samples were dehydrated and embedded in paraffin. Sections (3 mm-thickness) were cut and and immuflorescence were performed. The slides were incubated with the primary antibodies, rabbit anti-mice NLRP3 mAb (1:100, Cell Signaling), and rabbit anti-mice ASC polyclonal antibody (1:500, Abcam) overnight at 4°C. Goat anti-mouse Alexa Fluor 647 (Invitrogen) or goat anti-rabbit Alexa Fluor 594 (Invitrogen) was used as a secondary antibody. DAPI were used to stain nucleus. Images were acquired by an Axio Observer combined with an LSM 780 confocal device with 630× magnification (Carl Zeiss).

### Immunohistochemistry

Tissue sections from paraffin-embedded liver fragments obtained from infected or not mice were used for immunohistochemistry using NLRP3 (1:100 dilution; Cell Signaling) polyclonal antibody for *in situ* detection. Sequential immunoperoxidase labeling and erasing was then performed to determine additional markers after NLRP3 immune stain, using antibodies to ASC (1:100 dilution; Abcam). Before, the incubation with primary antibody, the slides were incubated with blocking solution, which is 1% of BSA and 0,5% of goat serum overnight. Then, the slides were incubated overnight with the primary anti-body. Next, the slides were washed and then mounted for immunoperoxidase polymer anti-mouse visualization system (SPD-125; Spring Bioscience, Biogen) and then with the chromogen substrate 3-amino-9-ethylcarbazole (AEC) peroxidase system kit (SK-4200; Vector Laboratories). Microphotographs after immunostaining of tissue slides were scanned on a VS120 Olympus microscope. After high-resolution scanning, slides with coverslips were removed in PBS and dehydrated through an ethanol gradient to 95% ethanol.

### Intravital microscopy

Confocal microscopy imaging was performed as previously described (Marques et al., 2015). Confocal images were acquired using an inverted Nikon Eclipse Ti microscope coupled to an A1R scanning head (Nikon) without modification. To visualize specific cell populations, prior to imaging, mice were intravenously administered a mixture or a single dose of Brilliant Violet 421 (BV421) anti-Ly6G fluorescent antibodies (0.5 μL/g, clone 1A8; Biolegend) for staining of neutrophils, anti-CD31 Fluorescein isothiocyanate (FITC) (0.75 μL/g, clone 390; eBioscience) for endothelial staining, anti-F480 phycoerythrin (PE) (0.25 μL/g, clone T45-2342; BD Biosciences) or allophycocyanin (APC) (0.25 μL/ g, BioLegend clone) for Kupffer cell staining and anti-CD45R (B220) Invitrogen Alexa Fluor 647 (Alexa 647) (0.25 μL/g, clone RA3-6B2; BD Biosciences) diluted in sterile saline. Digital quantification was performed by Volocity (6.3; PerkinElmer) and NIS-Elements (Nikon Instruments).

### Histological evaluation

Liver tissue samples were harvested, sectioned, and fixed with 10% formaldehyde and embedded in paraffin. Standard hematoxylin and eosin (H&E) stain were performed in 3 μm sections. Microphotographs after staining of tissue slides were scanned on a VS120 Olympus microscope and granulomas were measured using ImageJ. For the pathologist analysis, sections of 3-μm thick were stained with hematoxylin-eosin, and PAS (periodic acid-Schiff’s reactive). A pathologist, who was blinded to the experimental groups, analyzed the histological sections, and performed the semiquantitative analyses. For histomorphometry an image capture and analysis system consisting of a light microscope (Eclipse 800, Nikon) coupled to a digital camera (Evolution VR Cooled Color 13 bits, Media Cybernetics, USA) was used. The interface capture software used was Q-Capture 2.95.0, version 2.0.5 (Silicon Graphics Inc, USA); the high-resolution images 2048 X1536 pixel buffer were transmitted to a color LCD monitor in TIFF format and digitized. The images were captured after calibration of the appropriate color and contrast parameters and remained constant for each type of staining. Fifteen images from each animal were obtained using 40x objective lens, and evaluated. A histological scoring system was applied to verify the degree of liver injury and inflammation. For liver injury the histological parameters, vacuolization, ballooned cells, and steatosis were considered. The hepatic injury scoring is represented by the sum of steatosis (0–3), hepatocellular vacuolization (0-3), ballooned cells (0-2), poliploidia (0-3), and binucleation (0-3) scores, with the final score ranging between 0 and 14. Liver inflammation considered the sum of portal inflammation score (0-3) and lobular inflammation (number of granuloma/histological field).

### Statistical analysis

Statistical analysis were performed using the Prism 8 program (GraphPad Software, version 8.0). The values always present their medians and standard deviations obtained in each condition. All in vitro experiments were performed in triplicate and all in vivo experiments were performed using groups with at least 4 animals each. Values were analyzed using the Two-way ANOVA test or T test and 95% confidence intervals.

## Acknowledgments

We thank Maira Nakamura and Patricia Vendruscolo for the technical support and to members of the Zamboni lab for discussions and critical reading the manuscript. This work was funded by grants from the Center for Research on Inflammatory Diseases (CRID/CEPID/FAPESP, grant 2013/08216-2), Instituto Nacional de Ciência e Tecnologia em Vacinas (INCTV), CNPq (grant 401577/2014-7) and FAPESP (grants 2018/14398-0, 2019/11342-6, 2024/16091-0 and 2025/07812-8). D.S.Z. is a Research Fellow A from CNPq (grant 304945/2024-2).

## Author contributions

I.M.O designed and performed the experiments, analyzed the data, generated the figures, and wrote the manuscript. M.M.C, J.C.C.M. designed and performed the experiments, analyzed the data, generated the figures. A.L.B., P.S.C, B.O. designed and performed experiments and analyzed data. P.H.M., A.S, H.N. performed the bioinformatic analysis of single-cell RNA-seq and spatial transcriptomic datasets. T.S.R., A.M.B., L.L., S.S.B, A.J.R.H, L.L., performed experiments. A.M.S., R.T.G, A.K.C, G.B.M., C.H.S.C, provided reagents, designed experiments and discussed results and hypotheses. C.M.T. and A.T.F performed histopathological evaluation. D.S.Z. supervised the project, designed the experiments, helped with data interpretation, participated in the data analysis, and wrote the manuscript.

## Competing interests

The authors declare no competing interests.

## Notes

### Competing Interest Statement

The authors have declared no competing interest.

