## Supplemental Figures and Legends for "NLRP3 inflammasome signaling orchestrates the cellular and spatial architecture of hepatic granuloma and protective immunity"

### Supplementary Figures

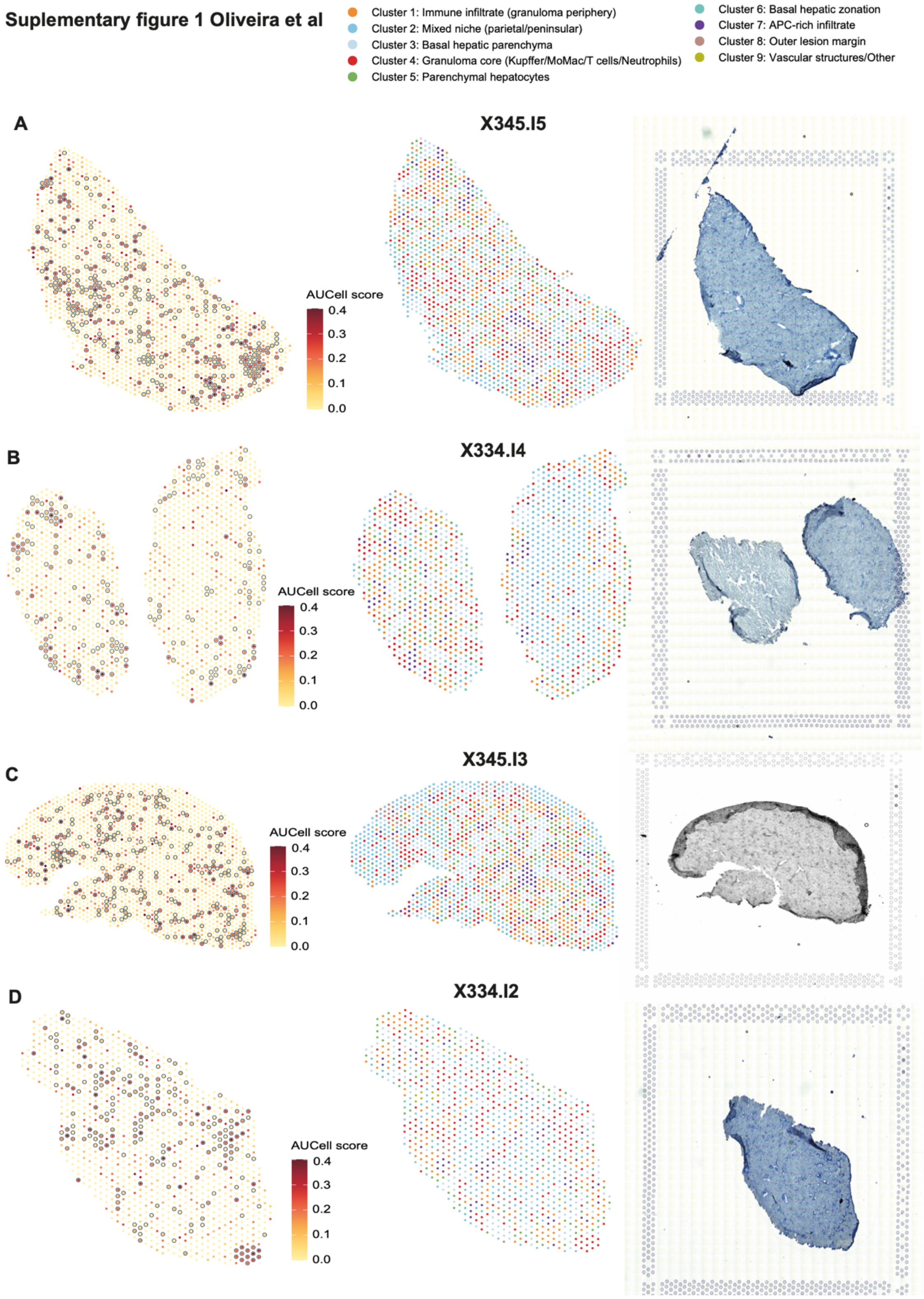

**Figure S1. Inflammasome-associated transcripts are spatially enriched in granuloma-associated regions during visceral leishmaniasis.** Spatial transcriptomic maps of samples X345.I5 (A), X334.I4 (B), X345.I3 (C) and X334.I2 (D). For each sample, the left panel shows the spatial distribution of the AUCell score for an inflammasome signature comprising *Nlrp3, Pycard, Casp1, Il1b, Il18* and *Gsdmd*. Spots assigned to cluster 4, corresponding to the granuloma core, are outlined. The middle panel shows the annotated spatial compartments, and the right panel shows the corresponding histological tissue section. Color scales indicate the inflammasome AUCell score and spatial-cluster identity.

#### Suplemmentary Figure 2 Oliveira et al

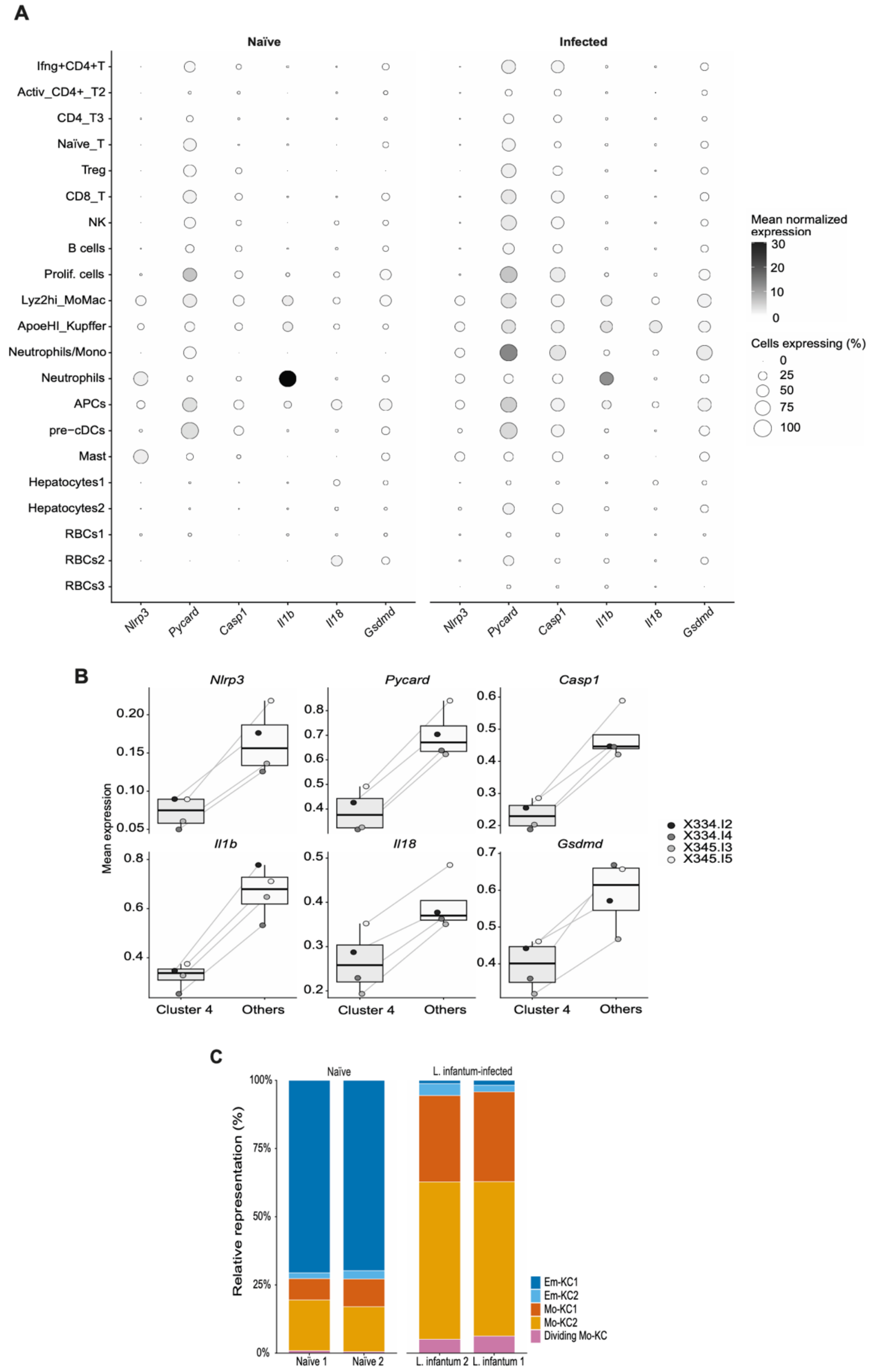

**Figure S2. Inflammasome-associated transcripts are enriched in hepatic macrophages during visceral leishmaniasis.** Dot plot showing the expression of *Nlrp3, Pycard, Casp1, Il1b, Il18* and *Gsdmd* across annotated liver cell populations from naïve and infected mice. Circle size represents the percentage of cells with detectable expression, whereas color indicates the mean expression of each gene, scaled as a *z* score (A). Mean spatial expression of the individual inflammasome-signature genes in cluster 4, corresponding to the granuloma core, and in the remaining spatial compartments. Each point represents one biological sample from the four spatial transcriptomic samples analyzed. Statistical significance was assessed using a permutation test (B). Relative composition of hepatic macrophage populations in two naïve and two *L. infantum*-infected samples. Stacked bars show the proportional representation of embryonically derived Kupffer cells, Em-KC1 and Em-KC2; monocyte-derived Kupffer cells, Mo-KC1 and Mo-KC2; and dividing Mo-KC cells (C). Each bar represents one biological sample.

**Supplementary figure 3 Oliveira et al**

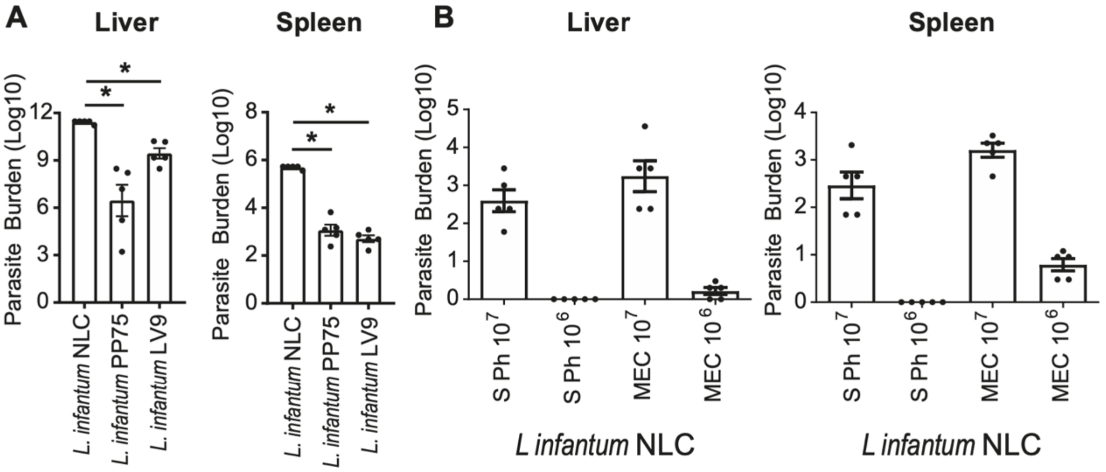


33

**Figure S3. The NLC strain of *L. infantum* efficiently establishes visceral infection in mice**. $10^7$ *L. infantum* Pp75, *L. infantum* NLC and *L. donovani* LV9 were injected by IP route in C57BL/6 mice. 3 weeks post infection, the animals were euthanized and the livers and spleen (A) were used to measure parasite burden. $10^6$ and $10^7$ *Leishmania spp* from SPh and MEC samples were injected into C57BL/6 mice by IP. After 3 weeks of infection, the animals were sacrificed and livers and spleens (B) removed to quantify the parasitic load by limiting dilution assay. *, $P < 0.05$.

**Supplementary figure 4 Oliveira et al**

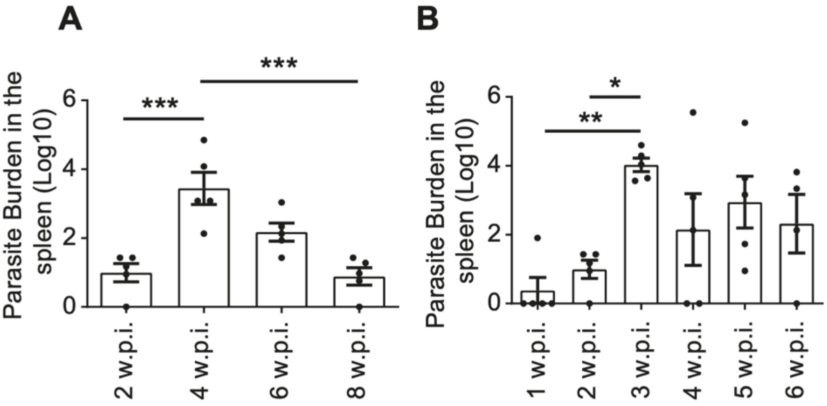

**Figure S4. The NLC strain of *L. infantum* efficiently establishes splenic infection in mice.** $10^7$ NLC *L. infantum* were purified from MEC and infected into C57BL/6 mice by IP route and followed for several weeks. After 2, 4, 6 and 8 weeks after the infection, a group of animals (n = 5) were sacrificed and the spleen (A) were removed to quantify parasite burden. Also, after 1, 2, 3, 4, 5 and 6 weeks after the infection, a group of animals (n = 5) were sacrificed and the spleens (B) were removed to quantify parasite burden. *, *P* p<0,05; **, *P* <0,005; ***, *P* <0,0005.

**Supplementary figure 5 Oliveira et al**

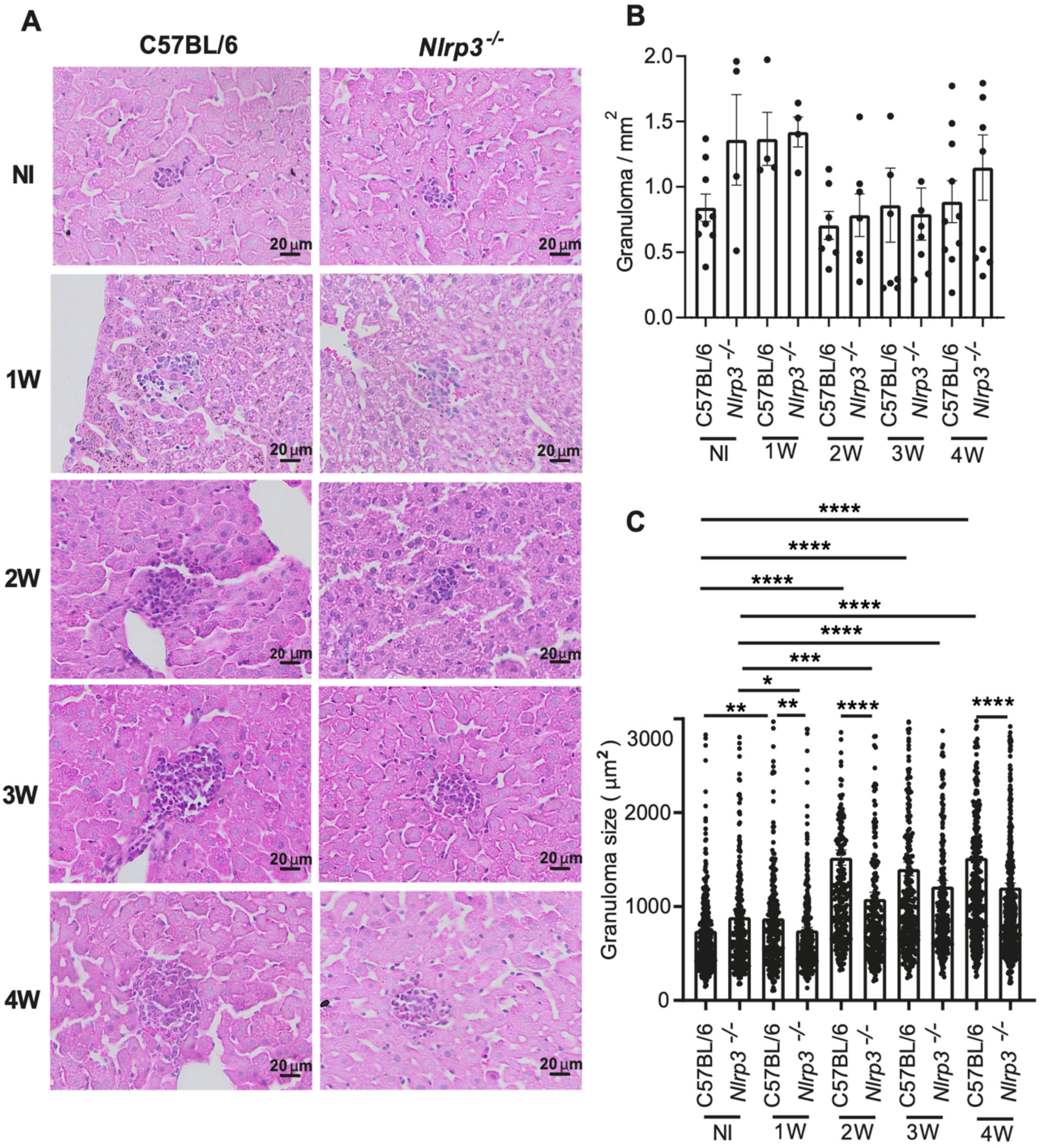


55

**Figure S5. The NLRP3 inflammasome promotes hepatic granuloma organization during *L. infantum* infection**. C57BL/6, *Nlrp3*$^{-/-}$ mice were infected with $10^7$ *L. infantum* MEC by IP route, the animals were euthanized and the livers were used to histological analysis. Livers from non-infected and infected C57BL/6 mice and *Nlrp3*$^{-/-}$ mice (A) were collected after 1, 2, 3 and 4 weeks pos infection. The number of granulomas per area were quantified using Image J (B). Granuloma size (C) were quantified using Image J. Each dot represents a granuloma from the animals shown in B. Set of 2 independent experiments were plotted together. *, $P$ <0.05; **, $P$ <0.005; ***, $P$ <0.0005; ****, $P$ <0.00005.

**Supplementary figure 6 Oliveira et al**

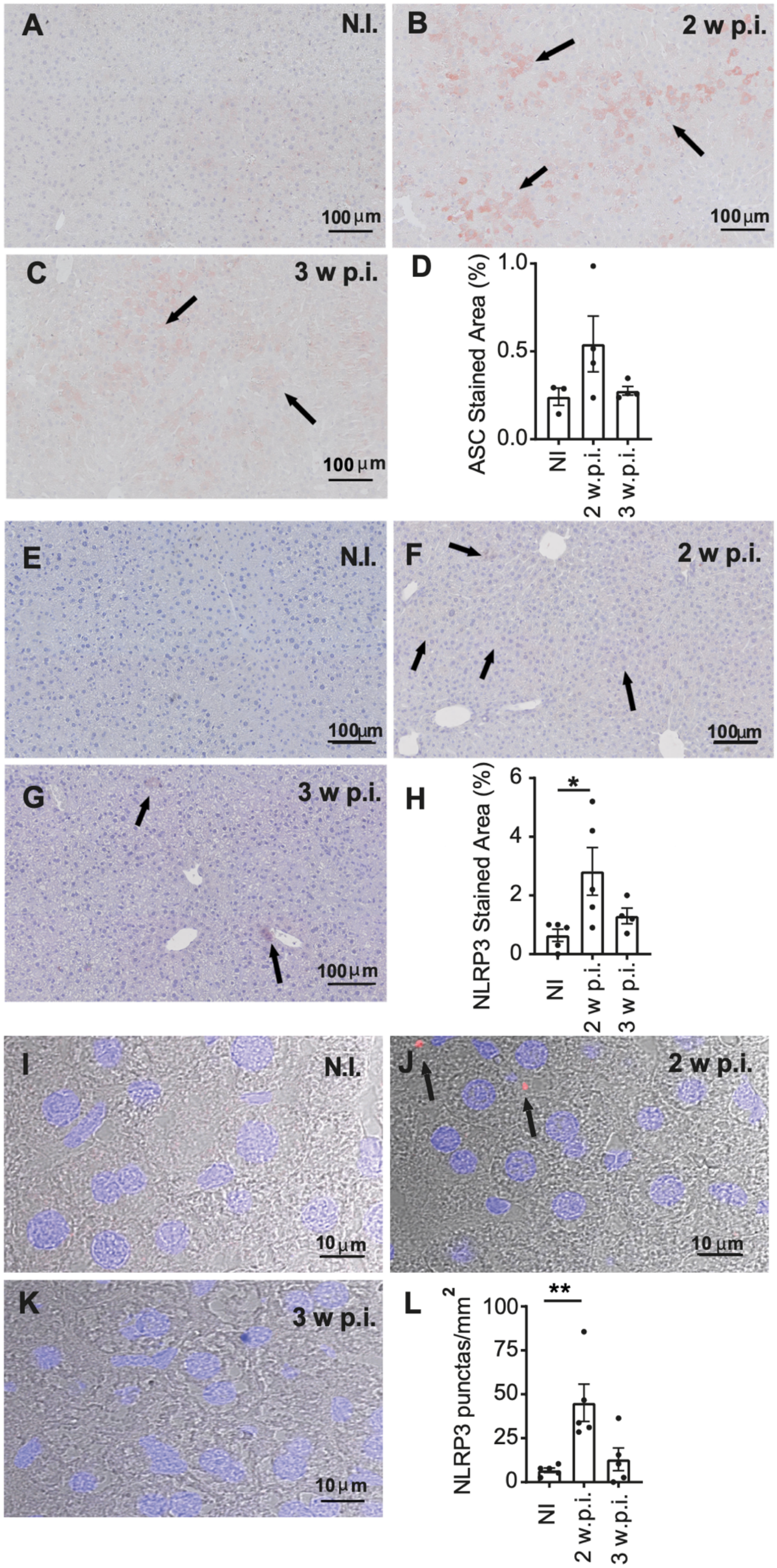


67

**Figure S6. NLRP3 and ASC are expressed and NLRP3 inflammasome is activated in the liver of *L. infantum*-infected mice.**

Livers from uninfected (NI) and infected C57BL/6 mice were collected 2 weeks (2 wpi) and 3 weeks (3 wpi) post-infection and stained for ASC (A-C) and NLRP3 (E-G). Representative immunohistochemical images of liver tissues. Quantification of ASC (D) and NLRP3 (H) expression in infected and uninfected livers. Representative immunofluorescence images of liver tissues from uninfected (NI) and infected C57BL/6 mice at 2 weeks (2 w.p.i) and 3 weeks (3 w.p.i) post-infection labeled with anti-NLRP3 (I-K) antibody. The number of NLRP3 puncta were quantified (L). *, $P <0.05$; **, $P <0,005$.

**Supplementary figure 7 Oliveira et al**

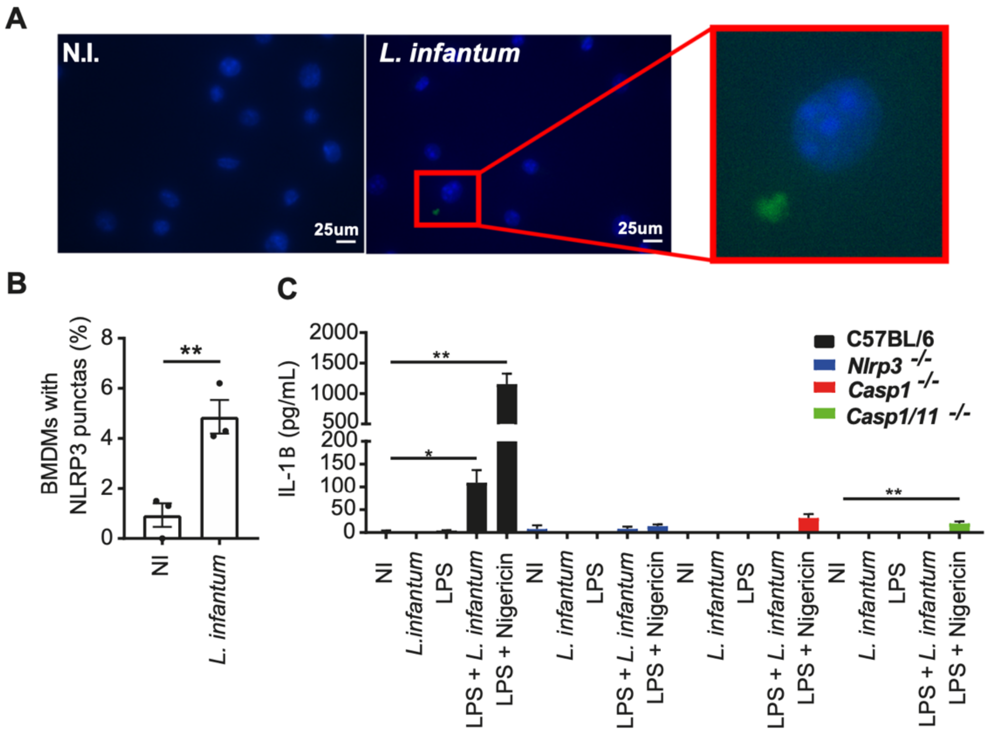


79

**Figure S7. *L. infantum* activates the NLRP3 inflammasome in BMDMs.**

Bone marrow-derived macrophages (BMDMs) were infected for 4 h with *L. infantum* NLC. After 24 h, non-infected and infected cells were treated with DAPI and anti-NLRP3, and NLRP3 puncta were quantified (A, B). BMDMs were pre-treated or not with 100ng/mL LPS for 4 h. Next, pre-treated or not BMDMs from C57BL/6, *Nlrp3*$^{-/-}$, *Casp1*$^{-/-}$ and *Casp1/11*$^{-/-}$ mice were or not infected with MOI10 from S Ph of NLC *L. infantum* culture for 24 h. 100 ng/mL LPS plus Nigericin was used as positive control. IL-1β production (C) was measured from supernatants. *, $P$<0,05; **, $P$<0,005; ***, $P$<0,0005.

**Supplementary figure 8 Oliveira et al**

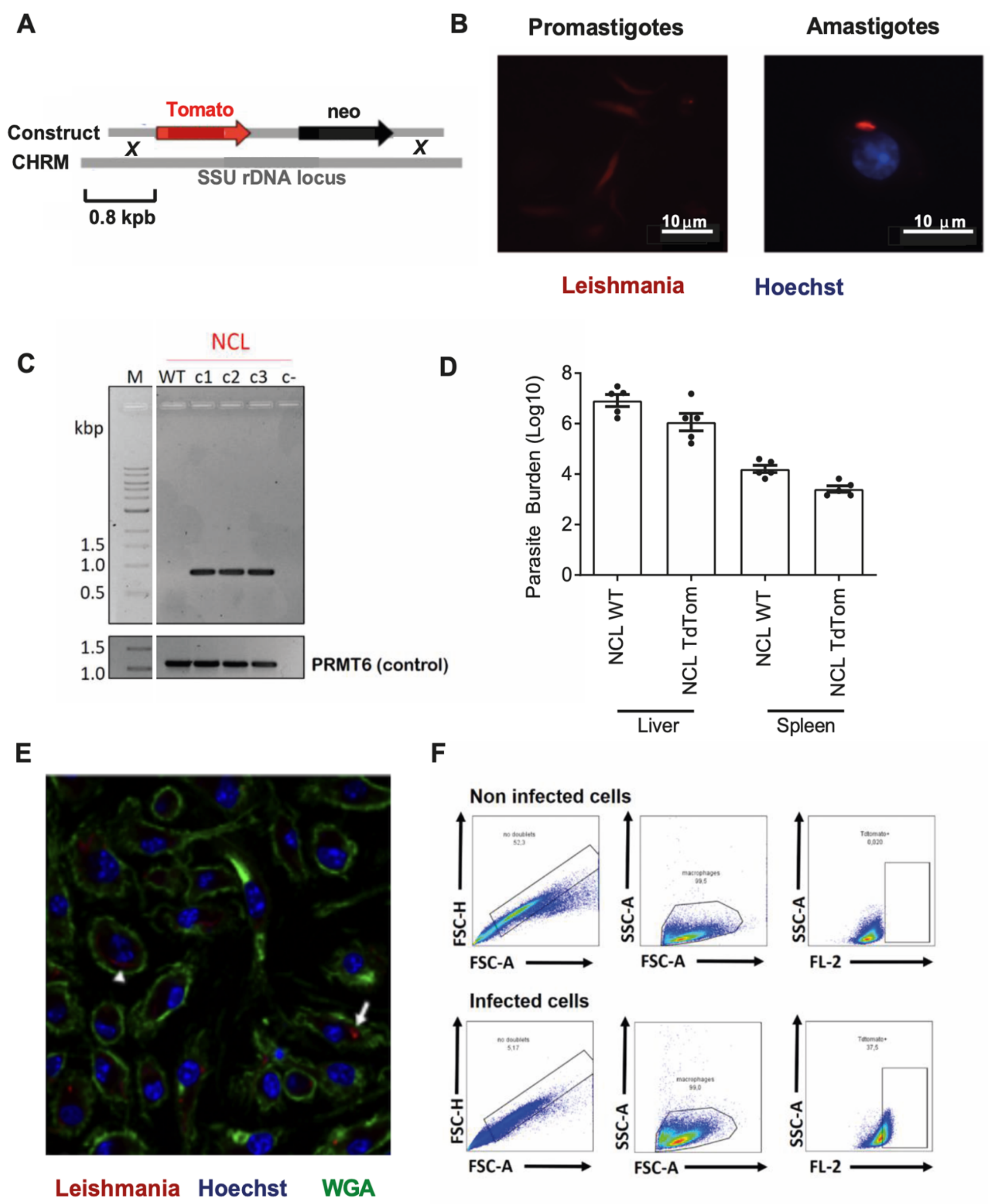

**Figure S8. Generation of a constitutively tdTomato-expressing *L. infantum* NLC strain**. *L. infantum* NLC parasites were transfected with a construct containing information to encode the fluorescent protein TdTomato, in addition to a gene that confers resistance to G418 (A). We observed the fluorescence of clone 3 in both promastigote and amastigote (B) parasites in the context of in vitro infection using BMDMs (macrophages with a blue-marked nucleus, using Hoechst, and red amastigote expressing tdTomato). The integration of the construction in the SSU locus was confirmed by PCR in three different clones (C). C57BL/6 mice were infected with $10^7$ MEC of TdTomato or WT *L. infantum* NLC strain for 3 weeks. Spleens and livers were collected and number of parasites was obtained using the limiting dilution assay (D). BMDMs were infected with MOI3 from MEC of NLC *L. infantum* (macrophages with a blue-marked nucleus, using Hoechst, green representing membrane marker and red amastigote expressing tdTomato) and fixed 2 h post infection. Arrow shows intracellular parasite and arrowhead indicates extracellular parasite (E). The fluorescence of the respective clones was compared using fluorescence microscopy, and the clone with the highest intensity was chosen to continue the experiments. The fluorescence emission was also evaluated by comparative flow cytometry (F) between uninfected macrophages and macrophages infected with the mutant strain. Data represented as mean ± DPM.

**Supplementary figure 9 Oliveira et al**

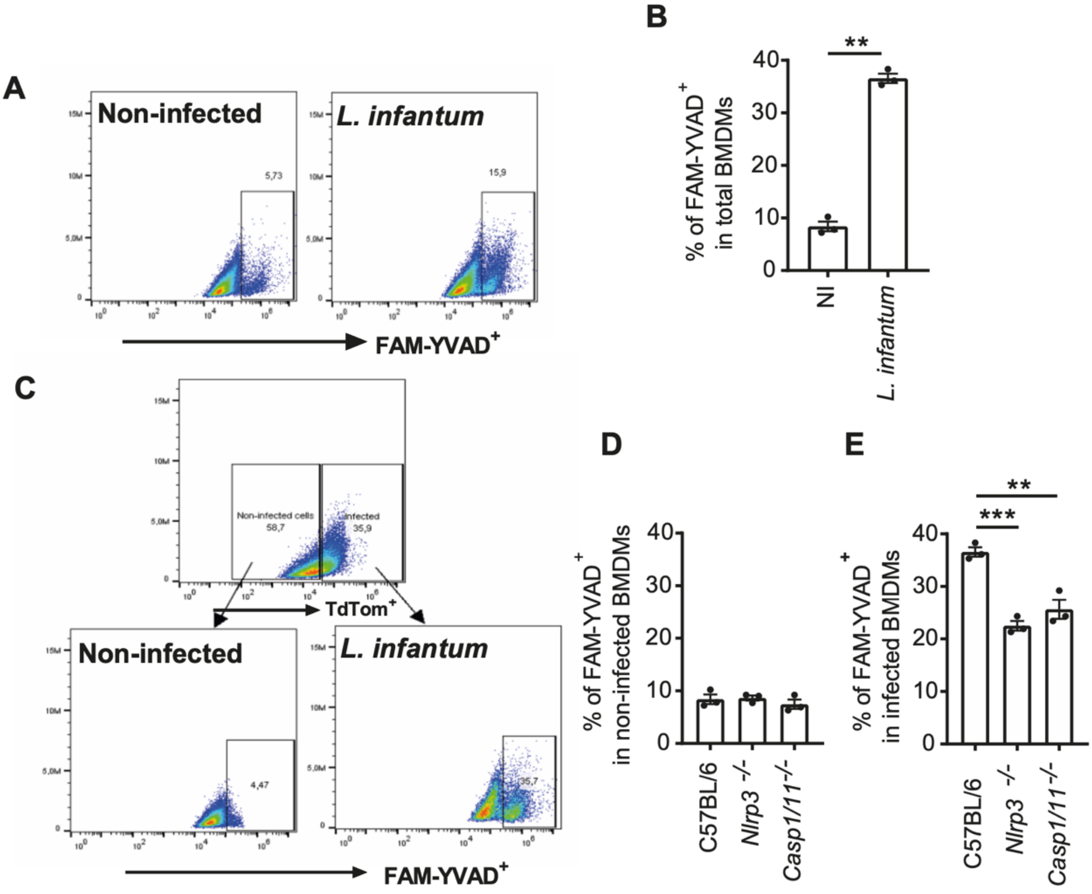


111

**Figure S9. *L. infantum* NLC infection induces caspase-1 activation in macrophages.** BMDMS were infected for 4 h with *L. infantum* NLC. BMDMs were pre-treated or not with 100ng/mL LPS for 4 h. Next, pre-treated or not BMDMs from C57BL/6, *Nlrp3*$^{-/-}$, *Casp1*$^{-/-}$ and *Casp1/11*$^{-/-}$ mice were or not infected with MOI10 from stationary phase *L. infantum* culture for 24 h. Caspase-1 activity production was measured from supernatants (A). Intracellular caspase-1 active was measured by flow cytometer using FLICA Caspase-1 Activity Kit in total cells (B). Gate in infected (C, E) and uninfected (D) was performed to determine the frequency of Caspase-1 activated in each population. Data represented as mean ± DPM; *=p<0,05; **=p<0,005; *** = p<0,0005.

**Supplementary figure 10 Oliveira et al**

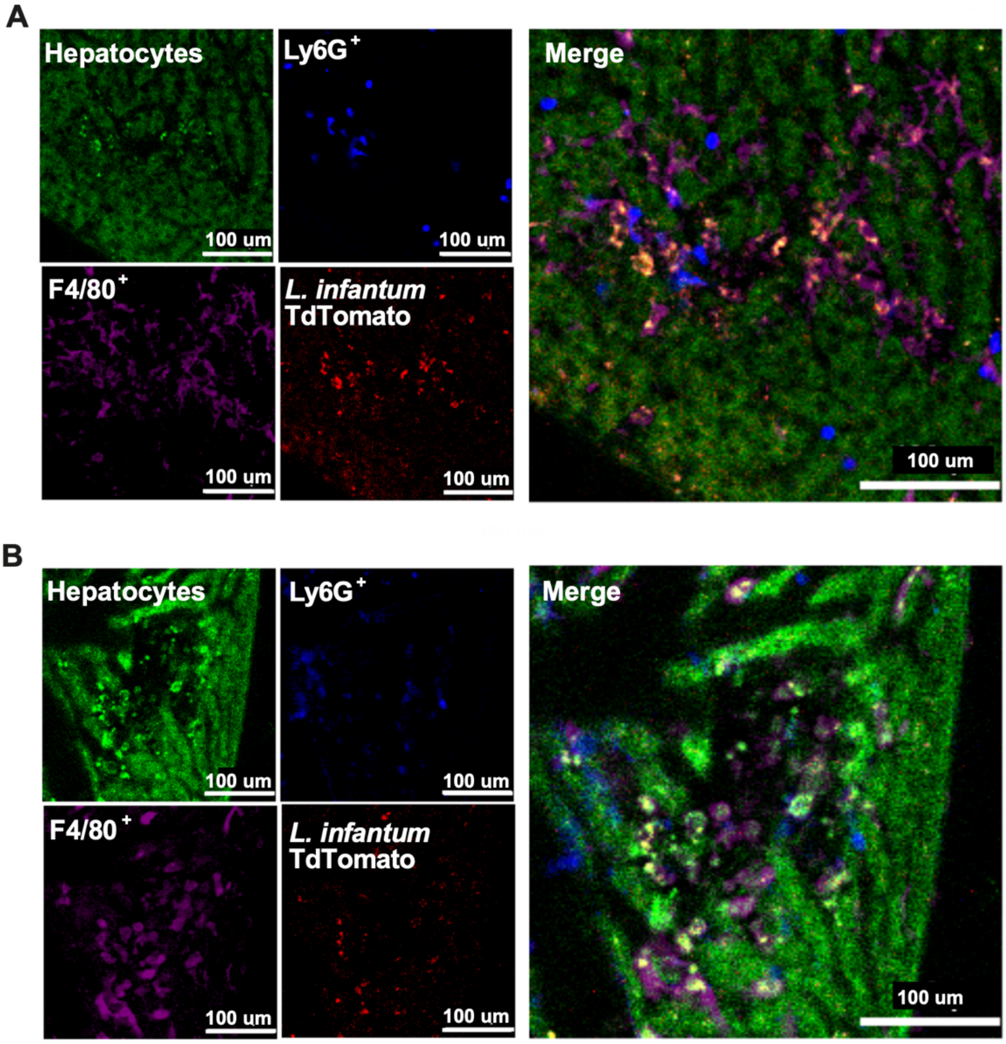


124

**Figure S10. *L. infantum* tdTomato-expressing parasites colocalize with Kupffer cells within hepatic granulomas of C57BL/6 mice**. C57BL/6 mice were infected with $10^7$ MEC of *L. infantum* TdTomato via IP route. Representative images of granulomas from C57BL/6 animals infected with *L. infantum* NLC TdTomato (A, B).

**Supplementary figure 11 Oliveira et al**

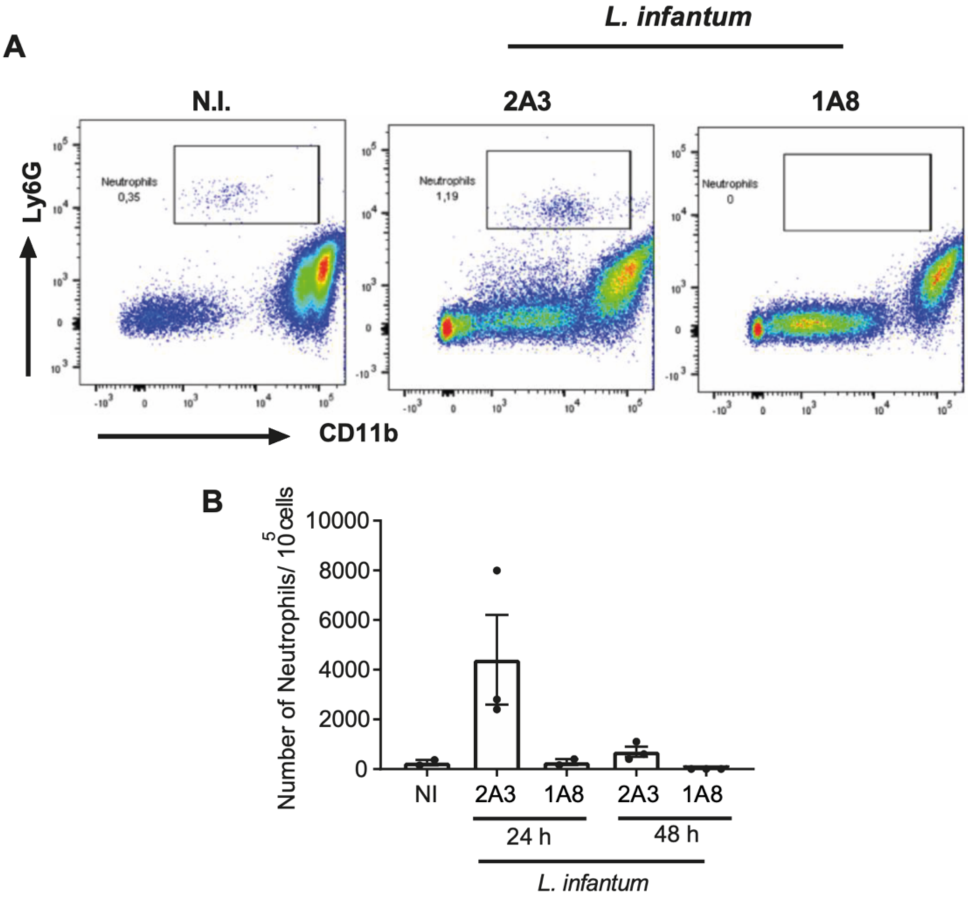

**Figure S11. Neutrophils are efficiently depleted using the anti-Ly6G antibody 1A8.** Neutrophils were depleted using 1 mg 1A8 treatment intraperitoneally. 1mg of 2A3 intraperitoneally were used as isotype control. Next, we injected $10^7$ metacyclic-enhanced cultures (MEC) of *L. infantum* NLC via IP.  Twenty-four and 48 h post infection, the numbers neutrophils, were determined by flow cytometry in the peritoneum (A, B). Data represented as mean ± SD. Each dot represents a single mouse.

**Supplementary figure 12 Oliveira et al**

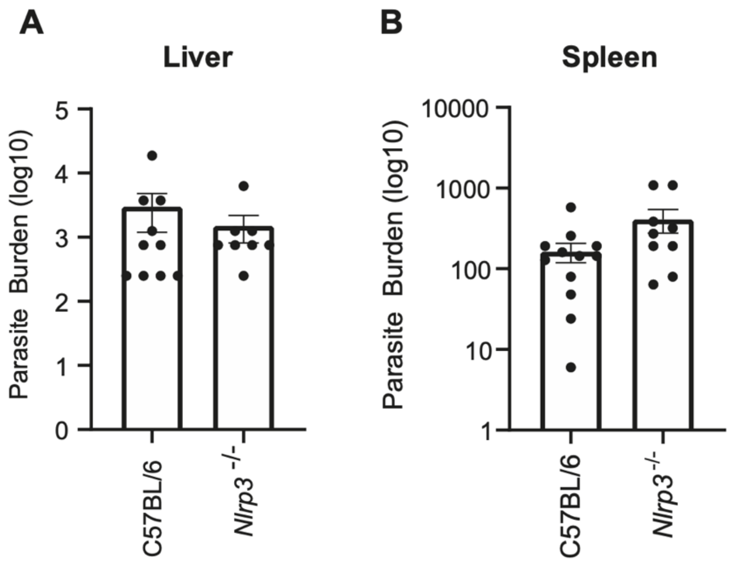

**Figure S12. NLRP3 deficiency does not affect parasite burden at 2 weeks after *L. infantum* infection**. C57BL/6 and *Nlrp3*$^{-/-}$ mice were infected with $10^7$ MEC of *L. infantum* NLC strain. After 2 weeks (2 w.p.i) post infection the liver (A) and spleen (B) were collected to access parasite burden.

**Supplementary figure 13 Oliveira et al**

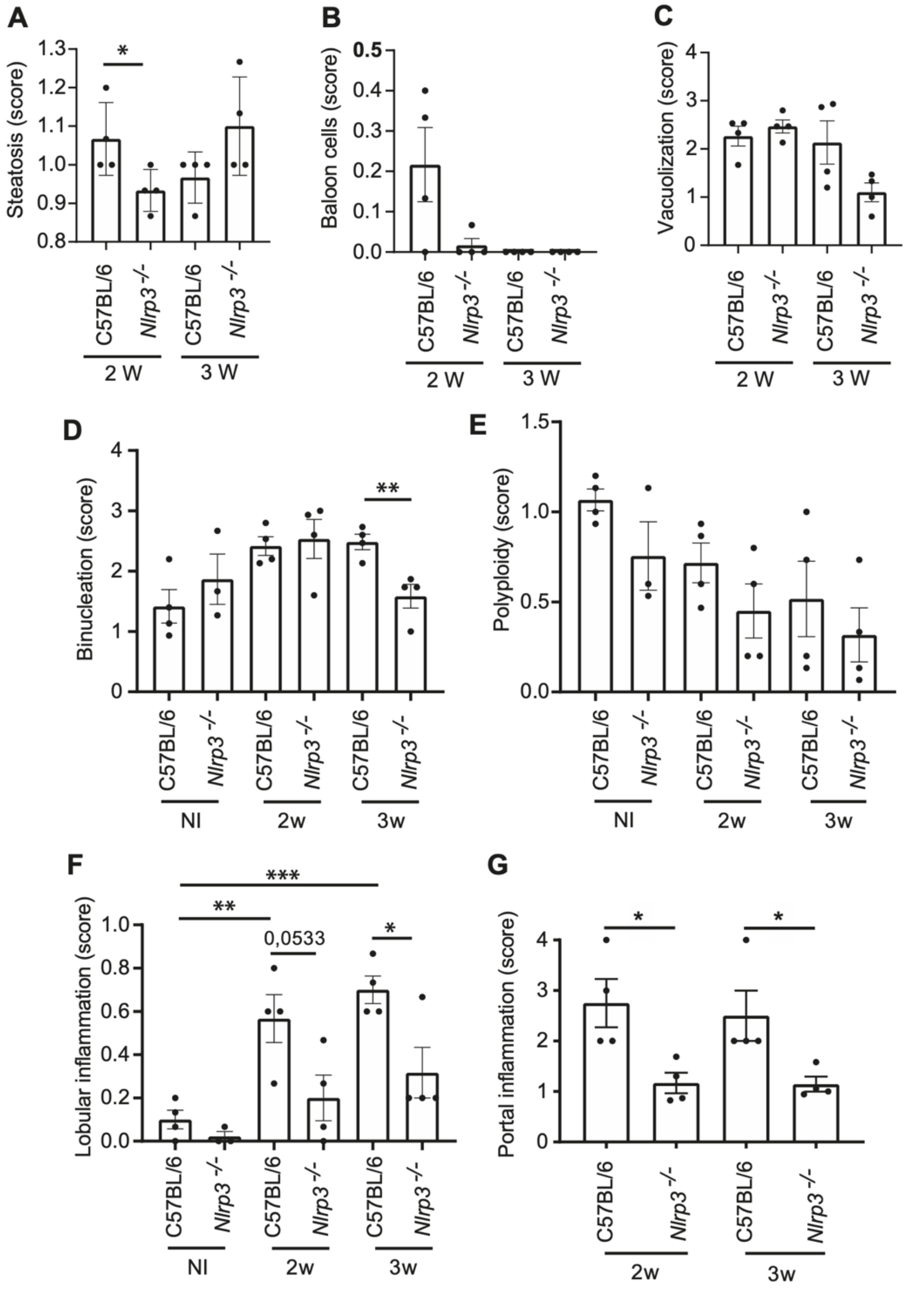

**Figure S13**. **Histological analysis of *L. infantum*-infected C57BL/6 and *Nlrp3*$^{-/-}$ mice.** Score of histological analyses of liver steatosis (A), ballon cells (B), vacuolization in cells (C), binucleation in liver cells (D), polyploidy (E), lobular inflammation (F) and portal inflammation (G). *, $P$ <0.05; **, $P$ <0.005; ***, $P$ <0.0005.
